# Coalescent Processes with Skewed Offspring Distributions and non-Equilibrium Demography

**DOI:** 10.1101/137497

**Authors:** Sebastian Matuszewski, Marcel E. Hildebrandt, Guillaume Achaz, Jeffrey D. Jensen

**Author notes:** contributed equally (Sebastian Matuszewski, Marcel E. Hildebrandt).

## Abstract

Non-equilibrium demography impacts coalescent genealogies leaving detectable, well-studied signatures of variation. However, similar genomic footprints are also expected under models of large reproductive skew, posing a serious problem when trying to make inference. Furthermore, current approaches consider only one of the two processes at a time, neglecting any genomic signal that could arise from their simultaneous effects, preventing the possibility of jointly inferring parameters relating to both offspring distribution and population history. Here, we develop an extended Moran model with exponential population growth, and demonstrate that the underlying ancestral process converges to a time-inhomogeneous psi-coalescent. However, by applying a non-linear change of time scale – analogous to the Kingman coalescent – we find that the ancestral process can be rescaled to its time-homogeneous analogue, allowing the process to be simulated quickly and efficiently. Furthermore, we derive analytical expressions for the expected site-frequency spectrum under the time-inhomogeneous psi-coalescent and develop an approximate-likelihood framework for the joint estimation of the coalescent and growth parameters. By means of extensive simulation, we demonstrate that both can be estimated accurately from whole-genome data. In addition, not accounting for demography can lead to serious biases in the inferred coalescent model, with broad implications for genomic studies ranging from ecology to conservation biology. Finally, we use our method to analyze sequence data from Japanese sardine populations and find evidence of high variation in individual reproductive success, but few signs of a recent demographic expansion.

---

The origins of the coalescent in the early 1970s mark a milestone for evolutionary theory (Kingman 2000). More than 45 years after Kingman formally proved the existence of the “*n*-coalescent” (Kingman 1982a,b,c), the so-called Kingman has gradually become the key theoretical tool to study the complex interplay of mutation, genetic drift, gene flow and selection. Closely linked to its underlying forward-in-time population model – e.g., the Wright-Fisher (WF; Fisher 1930; Wright 1931) and the Moran model (Moran 1958, 1962) – the Kingman coalescent has been used to derive expected levels of neutral variation, including the number of segregating sites *s*, the average number of pairwise differences *π*, and the distribution of the allele frequencies in a population ***η*** (i.e., the site frequency spectrum; SFS). In fact, these predictions not only apply to the WF and Moran model, but extend to a large class of Cannings exchangeable population models (Cannings 1974) that all converge to the Kingman coalescent in the ancestral limit (Möhle and Sagitov 2001). Furthermore, the Kingman coalescent forms the basis for many population genetic statistics – such as Tajima’s D (Tajima 1989), Fay and Wu’s H (Fay and Wu 2000) or more generally any SFS-based test statistic (Achaz 2009; Ferretti *et al.* 2010) – and subsequent inferences (Irwin *et al.* 2016) to detect deviations from the assumption of a neutrally evolving, constant-sized, panmictic population (Wakeley 2009).

While the Kingman coalescent has been shown to be robust to violations of its assumptions (Möhle 1998, 1999), such as constant population size, random mating, and non-overlapping generations, and has been extended to accommodate selection, migration, and population structure (Neuhauser and Krone 1997; Nordborg 1997; Wilkinson-Herbots 1998), it breaks down in the presence of skewed offspring distributions (Eldon and Wakeley 2006), strong positive selection (Neher and Hallatschek 2013), recurrent selective sweeps (Durrett and Schweinsberg 2004, 2005), and large sample sizes (Wakeley and Takahashi 2003; Bhaskar *et al.* 2014). In particular, all of these effects can cause more than two lineages to coalesce at a time, resulting in so-called multiple mergers. Hence, the underlying coalescent topology (i.e., the gene genealogy) is no longer represented by a bifurcating tree as in the “standard” Kingman case, but can take more complex tree shapes that can also feature several simultaneous mergers. Taking these points into account, a more general class of models, so-called multiple-merger coalescent (MMC) models, have been developed (e.g., Bolthausen and Sznitman 1998; Pitman 1999; Sagitov 1999; Schweinsberg 2000; Möhle and Sagitov 2001; reviewed in Tellier and Lemaire 2014), aiming to generalize the Kingman coalescent model (Wakeley 2013). As for the latter, these MMC models can often be derived from Moran models, generalized to allow multiple offspring per individual (Eldon and Wakeley 2006; Huillet and Möhle 2013; see also review of Irwin *et al.* 2016)).

Starting from such an extended Moran model, Eldon and Wakeley (2006) proved that the underlying ancestral process converges to a psi-coalescent (sometimes also called Dirac coalescent; Eldon *et al.* 2015), and that population genetic parameters inferred from genetic data from Pacific oysters (*Crassostrea gigas*) under this model vastly differ from those inferred assuming the Kingman coalescent. Their study – being the first to link MMC models to actual biological questions, molecular data and population genetic inferences – highlighted that high variation in individual reproductive success drastically affect both genealogical history and subsequent analyses; this has been observed in many marine organisms such Atlantic cod (*Gadus morhua*) and Japanese sardines (*Sardinops melanostictus*), but should also occur more generally in any species with type III survivorship curves that undergo so-called sweepstake-reproductive events (Hedgecock 1994; Hedgecock and Pudovkin 2011). Fundamentally, the problem is that an excess of low-frequency alleles (i.e., singletons), a ubiquitous characteristic of many marine species (Niwa *et al.* 2016), could be explained by either models of recent population growth or skewed offspring distributions when analyzed under the Kingman coalescent assuming neutrality which can result in serious mis-inference (e.g., a vast overestimation of population growth).

In developing a SFS-based maximum likelihood framework, Eldon *et al.* (2015) demonstrated that multiple merger coalescents and population growth can be distinguished from their genomic footprints in the higher-frequency classes of the SFS with high statistical power (see also Spence *et al.* 2016). However, there is currently neither a modelling framework that considers the genomic signal arising from the joint action of both reproductive skew and population growth nor is there any *a priori* reason to believe that the two could not act simultaneously.

Here, we develop an extension of the standard Moran model that accounts for both reproductive skewness and exponential population growth, and prove that its underlying ancestral process converges to a time-inhomogeneous psi-coalescent. By (non-linearly) rescaling branch lengths this process can – analogous to the Kingman coalescent (Griffiths and Tavaré 1998) – be transformed into its time-homogeneous analogue allowing efficient large-scale simulations. Furthermore, we derive analytical formulae for the expected site-frequency spectrum under the time-inhomogeneous psi-coalescent and develop an approximate-likelihood framework for the joint estimation of the coalescent and growth parameters. We then perform extensive validation of our inference framework on simulated data and show that both the coalescent parameter and the growth rate can be estimated accurately from whole-genome data. In addition, we demonstrate that when demography is not accounted for, the inferred coalescent model can be seriously biased with broad implications for genomic studies ranging from ecology to conservation biology (e.g., due to its effects on effective population size or diversity estimates). Finally, using our joint estimation method we re-analyze mtDNA from Japanese sardine (*Sardinops melanostictus*) populations and find evidence for considerable reproductive skew, but only limited support for a recent demographic expansion.

## Model and Methods

Here we will first present an extended, discrete-time Moran model (Moran 1958, 1962; Eldon and Wakeley 2006) with exponential population growth which will serve as the forward-intime population genetic model underlying the ancestral limit process. We will then give a brief overview of coalescent models with special focus on the psi-coalescent (Eldon and Wakeley 2006), before revisiting SFS-based maximum likelihood methods to infer coalescent parameters and population growth rates.

**Table 1.**
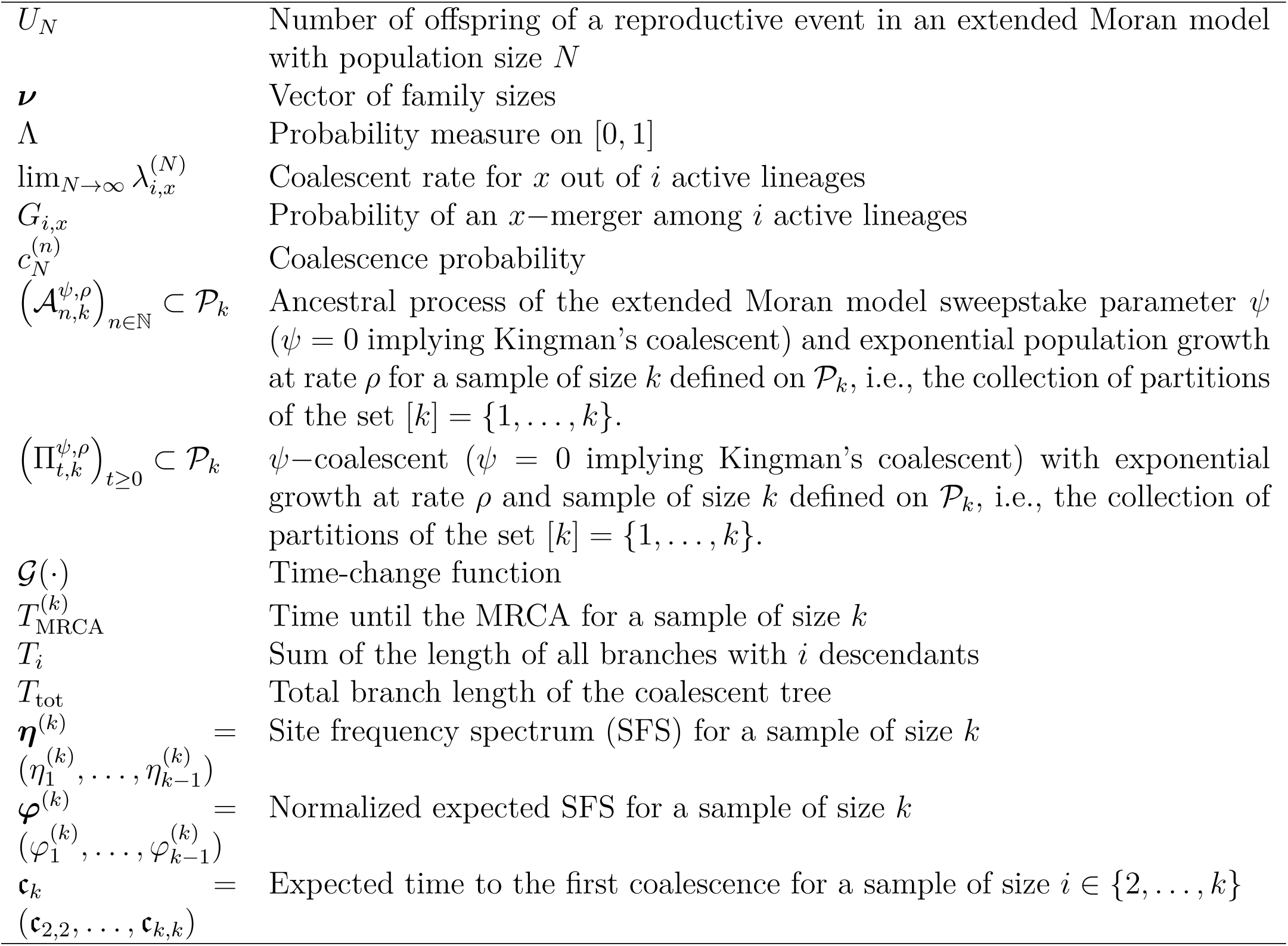
Summary of notation and definitions.

We consider the idealized, discrete-time model with variable population size shown generally in Figure 1. Furthermore, let *N*_*n*_ *∈* ℕ be the deterministic and time-dependent population size *n ∈* ℕ time steps in the past, where by definition *N* = *N*_0_ denotes the present population size. In particular, defining ***ν***(*n*) as the exchangeable vector of family sizes – with components *ν*_*i*_(*n*) indicating the number of descendants of the *i*^*th*^ individual – the (variable) population size can be expressed as

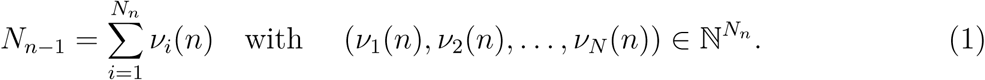

**Figure 1.**
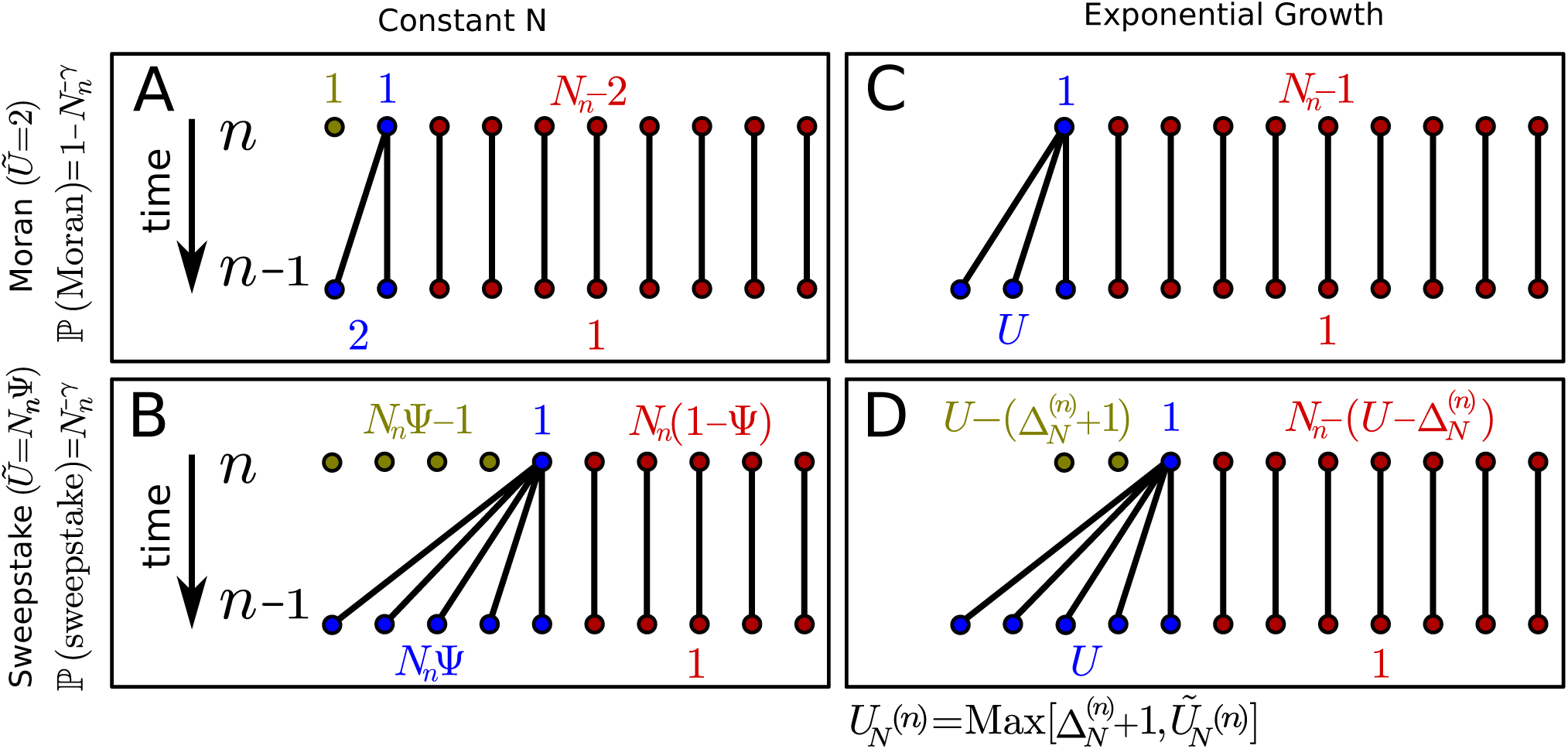
Illustration of the extend Moran model with exponential growth. Shown are the four different scenarios of population transition within a single discrete time step. **A** The population size remains constant and a single individual produces exactly two offspring (‘Moran-type’ reproductive event). **B** The population size remains constant and a single individual produces *ψN*_*n*_offspring (‘sweepstake’ reproductive event). **C** The population size increases by 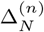 individuals and a single individual produces exactly 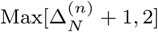 offspring. **D** The population size increases by 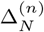 individuals and a single individual produces exactly 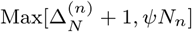 offspring. Note that *n* denotes the number of steps in the past, such that *n* = 0 denotes the present. An overview of the notation used in this model is given in Table 1.

Furthermore, we assume that the reproductive mechanism follows that of an extended Moran model (Eldon and Wakeley 2006; Huillet and Möhle 2013). In particular, as in the original Moran model, at any given point in time *n ∈*ℕ only a single individual reproduces and leaves *U*_*N*_ (*n*) offspring (including itself). Formally, the number of offspring can be written as a sequence of random variables (*U*_*N*_ (*n*))_*n∈*ℕ_ (where each *U*_*N*_ (*n*) is supported on 0, 1*, …, N*_*n–*1_), such that ***ν***(*n*) – up to reordering – is given by

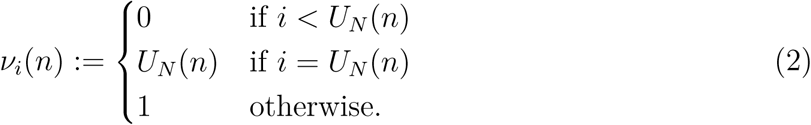

However, since population size varies over time, the sequence (*U*_*N*_ (*n*))_*n ∈*ℕ_ is generally not identically distributed. On a technical note though, we require that the (*U*_*n*_) are independently distributed which ensures that the corresponding backwards process satisfies the Markov property.

An illustration of our model and the four different scenarios for forming the next generation (i.e., within a single discrete time step) is shown in Figure 1. Generally we differentiate between two possible reproductive events: a classic ‘Moran-type’ reproductive event (Fig. 1**A**,**C**) and a ‘sweepstake’ reproductive event (Fig. 1**B**,**D**) occurring with probabilities 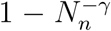 and 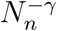, respectively. If the population size remains constant between consecutive generations (Fig. 1**A**,**B**), we re-obtain the extended Moran model introduced by Eldon and Wakeley (2006) in which a single randomly chosen individual either leaves exactly two offspring and replaces one randomly chosen individual (Moran-type) or replaces a fixed proportion *ψ ∈* (0, 1] of the population (of size *N*_*n*_). Note that throughout, without loss of generality, we assume that *N*_*n*_*ψ* is integer-valued. In both reproductive scenarios the remaining individuals persist. However, if the population size increases between consecutive generations (Fig. 1**C**,**D**), the reproductive mechanism needs to be adjusted accordingly. Let

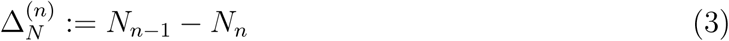

denote the increment in population size between two consecutive time points. Then the number of offspring at time *n* is given by

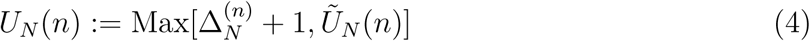

where 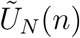 denotes number of offspring for the constant-size population. Thus, independent of the type of reproductive event – i.e., Moran-type or sweepstake – and in the spirit of the original Moran model, additional individuals are always assigned to be offspring of the single reproducing individual of the previous generation.

Following Eldon and Wakeley (2006), the distribution of the number of offspring ℙ (*U*_*N*_ (*n*) = *u*) can be written as

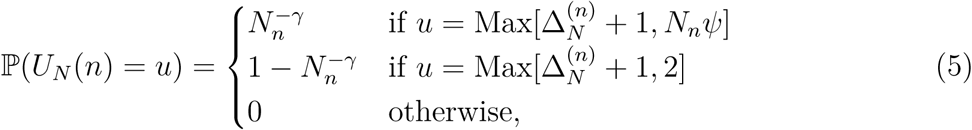

for some *γ >* 0 that – for a given fixed population size – determines the probability of a sweepstake reproductive event. We will here only consider the case where 1 *< γ <* 2 such that sweepstake events happen frequently enough that the ancestral process will be characterized by multiple mergers, and that all coalescent events are due to sweepstake reproductive events, but not so frequently that the population is devoid of genetic variation (Eldon and Wakeley 2006). Note that while the numbers of offspring and replaced individuals are no longer (necessarily) equal when the population size increases, the general reproductive mechanism remains unaltered.

Throughout the paper and following Griffiths and Tavaré (1994), we will assume that the population is growing exponentially over time at rate ϱ, and in particular that the population size, *n* steps in the past, is given by

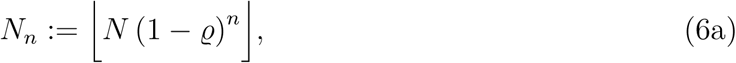

with

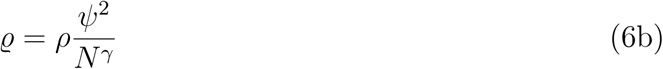

if the ancestral process is dominated by sweepstake events (i.e., if 1 *< γ <* 2), or

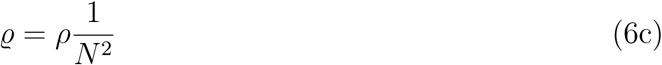

if Moran-type reproductive events dominate (i.e., if *γ >* 2), and the growth rate *ρ* is measured in units of the corresponding coalescent time. A discussion and details about the derivation of the coalescent-time scaling are given below in the *Derivation of the ancestral limit process* section.

### Multiple merger coalescents: The Psi-coalescent

The most general class of coalescent processes that allows for multiple lineages to coalesce per coalescent event (but not for multiple coalescent events at the same time) is the so-called Λ-coalescent. These processes are partition-valued exchangeable stochastic processes defined by a finite measure Λ on the [0, 1] interval (Donnelly and Kurtz 1999; Pitman 1999; Sagitov 1999). In particular, the rate at which *x* out of *i* active lineages merge is given by

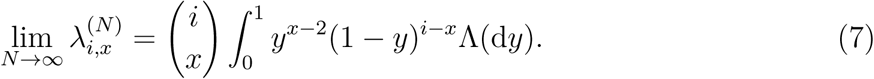

Special instances of the Λ-coalescent are Kingman’s coalescent (Kingman 1982a,b) with

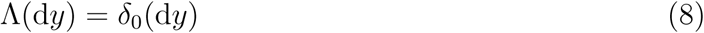

and the psi-coalescent (Eldon and Wakeley 2006) with

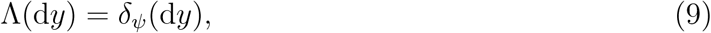

where the measure Λ is entirely concentrated at 0 and *ψ*, respectively.

Under a (constant-size) extended Moran model as proposed by Eldon and Wakeley (2006) (corresponding to Fig. 1**A**,**B**), the scaled coalescence rates of the ancestral process become

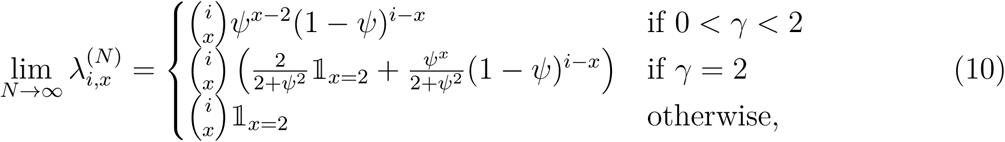

where 𝕝_*x*=2_ denotes the indicator function that is 1 if *x* = 2 and 0 otherwise. Accordingly, the corresponding rate matrix of the ancestral process ***Q***_*ψ*_*∈* ℝ^*k×k*^ with sample size *k* is given by

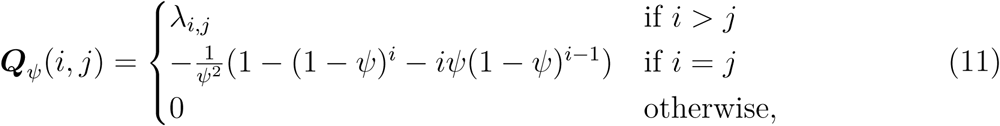

where *j* = *i – x –* 1. Note that the diagonal entries of ***Q***_*ψ*_ (*i, j*) (i.e., when *i* = *j*) is given by the (negative) sum over all coalescent rates, i.e., 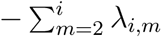, which evaluates to the closed-form representation given in the second line of Equation 11.

In particular, in the boundary case *ψ* = 0 we recover the rate matrix under the Kingman coalescent as

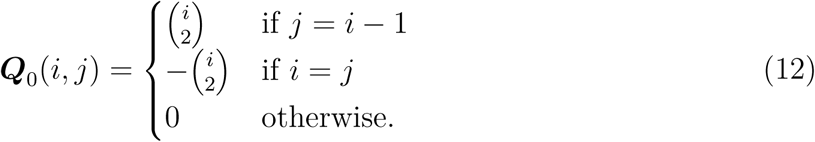

Note that in the infinite population size limit *γ* defines the time scale of the ancestral process. In particular, if 0 *< γ <* 2 all coalescence events are due to sweepstake reproductive events, whereas sweepstake events do not happen frequently enough if *γ >* 2, such that all (2-)mergers are due to Moran-type reproductive events. Moreover, in the latter case the ancestral process of the Moran model can be accurately described by the Kingman coalescent (when scaled appropriately). Note that for the special case *γ* = 2 both reproductive events happen on the same time scale (Eldon and Wakeley 2006).

### SFS-based maximum likelihood inference

In order to infer the coalescent model and its associated coalescent parameter and to (separately) estimate the demographic history of the population, Eldon *et al.* (2015) recently derived an (approximate) maximum likelihood framework based on the SFS (see also Birkner and Blath 2008 and Koskela *et al.* 2015 for alternative inference approaches based on a full likelihood framework and approximate conditional sampling distributions, respectively).

In the following we will give a concise overview of their approach which forms the basis for the joint inference of coalescent parameters and population growth rates.

First, let *k* denote the number of sampled (haploid) individuals (i.e., the number of leaves in the coalescent tree). Furthermore, let 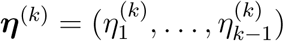 denote the number of segregating sites with derived allele count of *i* = 1,…,*k*–1 of all sampled individuals (i.e., the SFS), and let 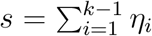 be the total number of segregating sites. Provided that *s >* 0 we define the normalized expected SFS 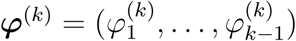 as

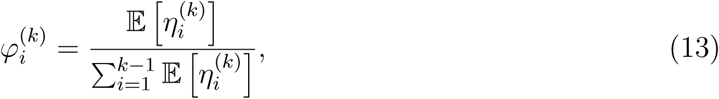

which – given a coalescent model 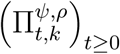 and assuming the infinite-sites model (Watterson 1975) – can be interpreted as the probability that a mutation appears *i* times in a sample of size *k* (Eldon *et al.* 2015). Furthermore, note that 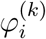 is a function of 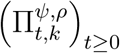 (i.e., of the coalescent process and the demographic population history), but unlike 𝔼[***η***^(*k*)^] is not a function of the mutation rate, and should provide a good first-order approximation of the expected SFS as long as the sample size and the mutation rate are not too small (Eldon *et al.* 2015).

Then the likelihood function 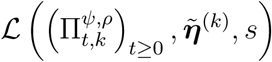 for the observed frequency spectrum 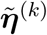 and given coalescent model 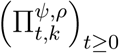 is given by

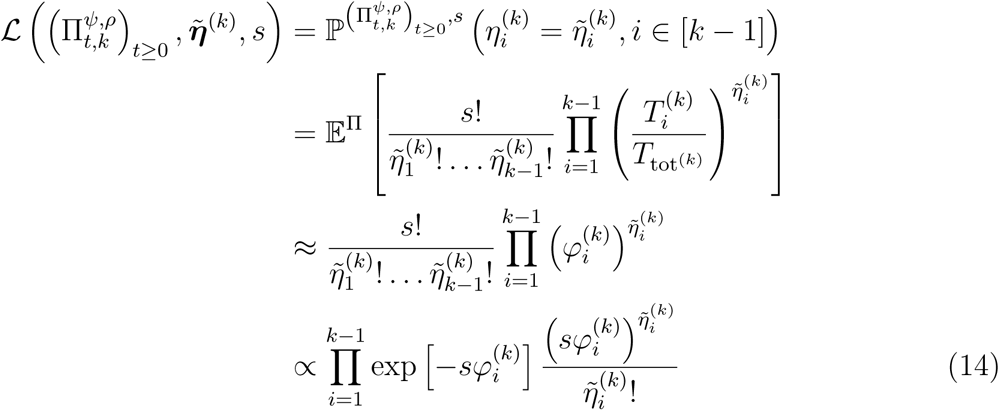

(Eldon *et al.* 2015). Note that in the third line we approximated 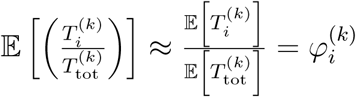. In fact, Bhaskar *et al.* (2015) recently used a Poisson random field approximation to derive an analogous, structurally identical likelihood function for estimating demographic parameters under the Kingman coalescent. Notably though, their approximation assumes that the underlying coalescent tree is independent at each site, under which condition Equation 14 is exact.

As an alternative to the likelihood approach, we followed Eldon *et al.* (2015) and also implemented a minimal-distance statistic approach where

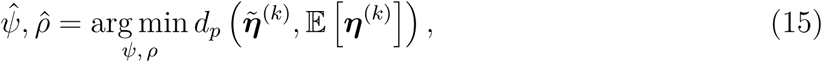

where *d*_*p*_ is some metric on ℝ^*p*–1^ calculated between the observed and the expected SFS under the generating coalescent process.

Note though that both the likelihood and the distance-based approach require expressions for the normalized expected SFS *φ*^(*k*)^. Instead of performing Monte Carlo simulations to obtain these quantities we adapted an approach recently proposed by Spence *et al.* (2016), who derived analytical formulas for the expected SFS under a given (general) coalescent model 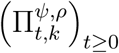 and an intensity measure ξ(*t*) : R_*≥*0_ *→* R_*>*0_. In particular, the authors showed that

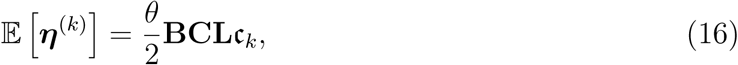

where **B** *∈* ℝ ^*k*–1*×k*–1^ and **C** *∈* ℝ ^*k*–1*×k*–1^ are both Λ independent (and thus easy to calculate) matrices, **L***∈* ℝ^*k*–1*×k*–1^ is a Λ dependent lower triangular matrix that depends on the rate matrix **Q** and its spectral decomposition, *θ* is the population-scaled mutation rate, and c_*k*_ = (c_2,2_, …, c_*k,k*_) denotes the expected time to the first coalescence for a sample of size *i ∈* {2*, …, k*}. Importantly, the time-inhomogeneity of the underlying coalescent process only enters through the first coalescence times c_*k*_. For example, the first coalescence times for the Kingman coalescent with an exponentially growing population are given by

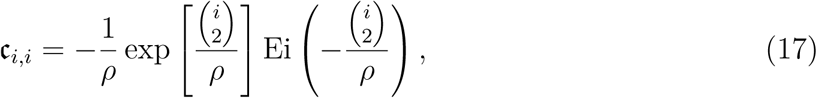

where 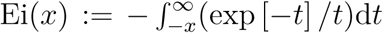 denotes the exponential integral (Polanski *et al.* 2003; Polanski and Kimmel 2003; Bhaskar *et al.* 2015). Finally, plugging Equation 16 into Equation 13 leads to

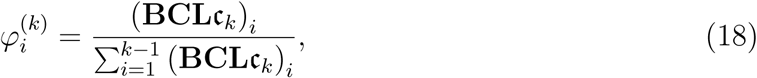

highlighting that *θ* cancels and that the likelihood function (eq. 14) is independent of the mutation rate.

To obtain the coalescent parameter *ψ* and population growth rate *ρ* that maximize the likelihood function (eq. 14) or respectively minimize the distance function (eq. 15), we used a grid search procedure over an equally-spaced two-dimensional grid with *ψ*_grid_ = {0, 0.01*, …,* 1} and *ρ*_grid_ = {0, 1*, …,* 1024}, and evaluated the value of the likelihood respectively distance function at each grid point.

### Data availability

The empirical raw data used have been downloaded from GenBank (accession numbers LC031518 – LC031623; data from Niwa *et al.* 2016). The empirical SFS can be downloaded from Supporting Information (**Supporting Files**: File E1). The simulation program and the inference program were written in C++ and are available upon request.

## Results and Discussion

The aim of this work is to derive the ancestral process for an exponentially expanding population that undergoes sweepstake reproductive events. We first derive the time-inhomogeneous Markovian ancestral process that underlies the extended Moran model, and show that, analogous to the Kingman coalescent, it can be described by a time-homogeneous Markov chain on a non-linear time scale. In particular, we derive the coalescent rates and the time-change function, and prove convergence to a Λ –coalescent with Dirac measure at *ψ*. Detailed derivations of the results, which in the main text have been abbreviated to keep formulas concise, can be found in Supporting Information (**Detailed derivation of results**). On the basis of these results, we derive a maximum likelihood inference framework for the joint inference of the coalescent parameter and the population growth rate, and assess its accuracy and performance through large-scale simulations. Furthermore, we quantify the bias of coalescent and population growth parameter estimates when mistakenly neglecting population demography or reproductive skew. Finally, we apply our approach to mtDNA from Japanese sardine (*Sardinops melanostictus*) populations where patterns of sequence variation were shown to be more consistent with sole influence from sweepstake reproductive events, again highlighting the potential mis-inference of growth if reproductive skew is not properly accounted for (Niwa *et al.* 2016; Grant *et al.* 2016).

### Derivation of the ancestral limit process

Unlike in the case of a constant-size population, the sequence of the number of offspring (*U*_*N*_ (*n*))_*n∈*N_ changes along with the (time-dependent) population size. Thus, the ancestral process is characterized by an inhomogeneous Markov chain with transition probabilities

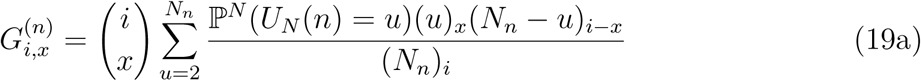

where (*z*)_*j*_ is the descending factorial, *z*(*z–*1) *…* (*z–j* + 1) with (*z*)_0_ = 1, and ℙ^*N*^ denotes the rescaled distribution of *U*_*N*_ given by

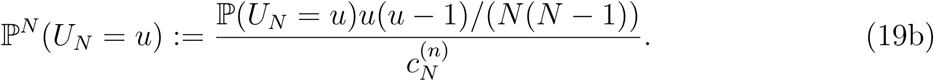

Note that ℙ^*N*^ is scaled by the time-dependent coalescence probability 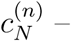 which scales the unit of time in the limit process such that it is equal to *G*_2,2_ steps in the discrete-time model and thus serves as the “natural” time scale for the corresponding ancestral process – defined as

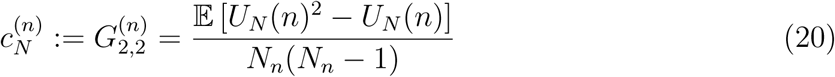

for all *n ∈* ℕ.

Plugging Equation 5 into Equations 19a and 20, and using Equation 4 then yields

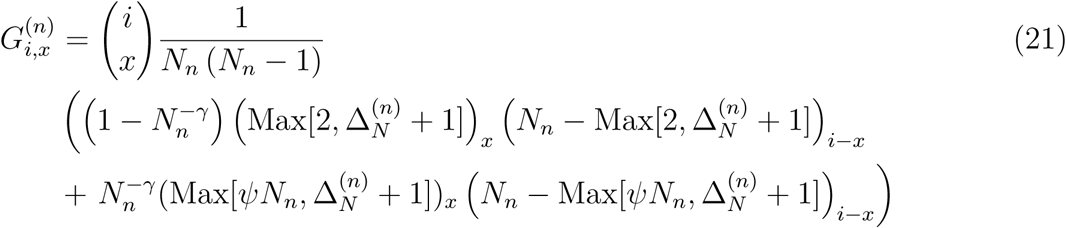

and

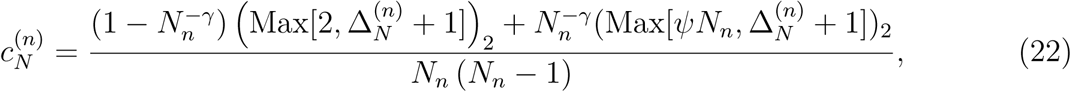

respectively. Note that Equation 22 is the weighted sum of the number of offspring for the two different reproductive events. Furthermore, taking the limit *N → ∞* in Equation 3

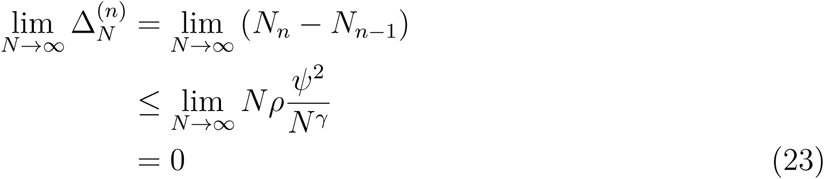

shows that 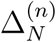 is bounded for all *n ∈* ℕ under the exponential growth model and thus allows dropping of the maxima condition in Equations 21 and 22. Furthermore, for sufficiently large *N* Equation 22 becomes

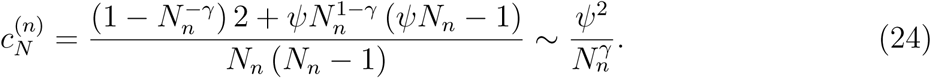

To prove that the time-scaled ancestral process of the underlying extended Moran model converges to a continuous-time Markov chain as the initial population size approaches infinity, we apply *Theorem 2.2* in Möhle (2002), which requires the following definitions: First, consider a step function *F*_*N*_ : [0*, ∞*) *→* [0*, ∞*) given by

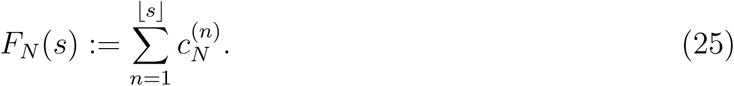

Furthermore, let 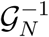 denote a modification of the right-continuous inverse of *F*_*N*_

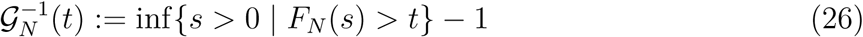

which will constitute the time-change function in the following. Since by assumption lim_*s→ ∞*_*F*_*N*_ (*s*) = *∞* it follows that 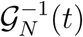 is finite for all *t ∈* [0*, ∞*). Finally, *Theorem 2.2* (Möhle 2002) requires that for all *t ∈* [0*, ∞*)

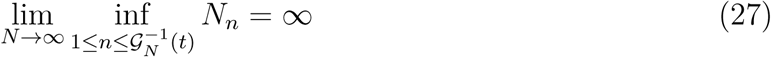

and

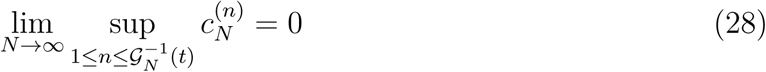

holds, i.e., that – on the new time scale – the population size remains large while the coalescent probabilities become small.

Then, let 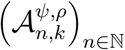 denote the ancestral process of the extended Moran model with exponential growth (see Model and Methods) and let *ϕ* and *ξ* denote two partitions of [*k*] ξ ⊂ *ϕ* with of size *a* and *b* = *b*_1_ + … + *b*_*a*_ (where *b*_1_ ≥ *b*_1_ *b*_2_ ≥ *b*_a_ ≥ 1), respectively. The transition probability of 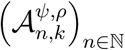 at time *n ∈* ℕ is given by

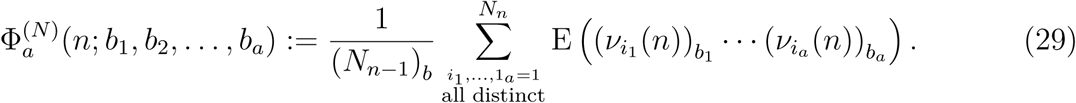

Thus, for the extended Moran model with exponential growth *Theorem 2.2* in Möhle (2002) states:

#### Theorem 1

(*Theorem 2.2*; Möehle 2002). *Assume that Equations (27) and (28) hold and for all t ∈* ℝ_*>*0_ *the limit*

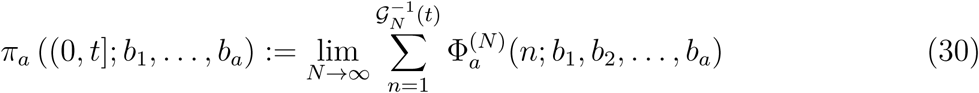

*exists. Then for each sample size k ∈* ℕ *the ancestral process 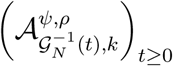 converges as N tends to infinity to a time-continuous and in general a time-inhomogeneous Markov chain 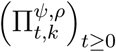.*

Note though that in its general form Theorem 1 was derived for any generic Cannings model as well as any kind of population size change (Möhle 2002).

We will now derive our first main result and show that the ancestral limiting process 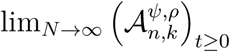 converges to a Λ *– k–*coalescent on a non-linear time scale. First, we derive the time-change function 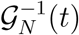 for the ancestral process by considering the step function (eq. 25)

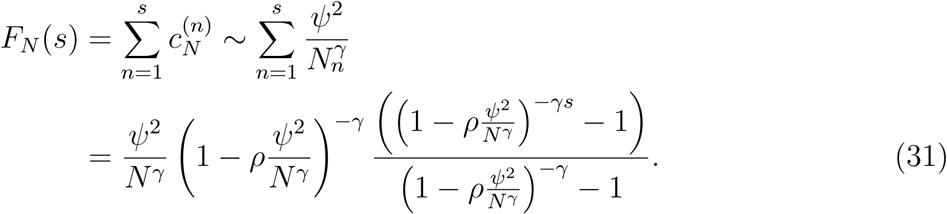

Solving for *s* then gives

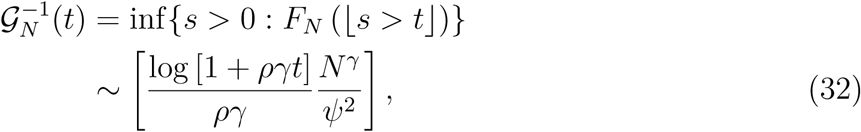

where we have used 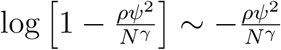 for sufficiently large *N*. In particular, we have

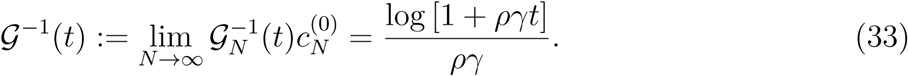

Furthermore, Equations 27 and 28 hold since

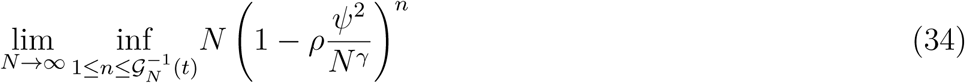

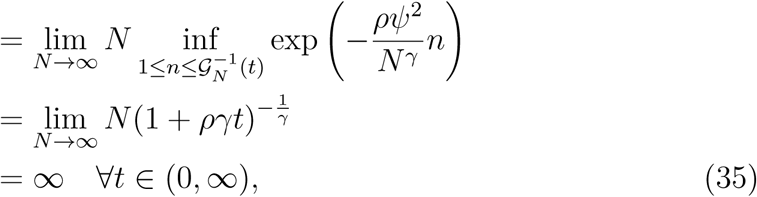

and by the same reasoning

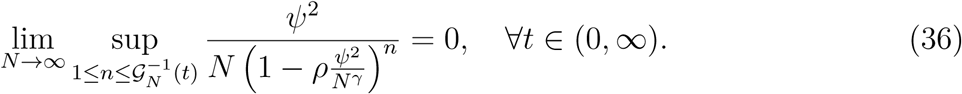

Finally, to show that Equation 30 holds, we first note that

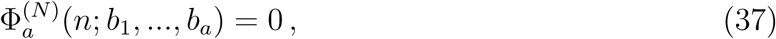

and for *a ≥* 2 and there are two indices 1 ≤*i < j* ≤ *a* with *b*_*i*_*, b*_*j*_ *≥* 2

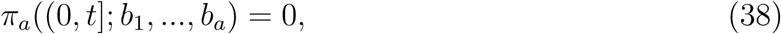

since the extended Moran model does not allow for more than one reproductive event at a time.

Thus, 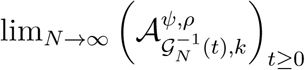 is well defined and does not feature any simultaneous coalescent events, implying that the limiting process must be a (possibly time-inhomogeneous) Λ – *k–*coalescent. Further, for *a* = 1,

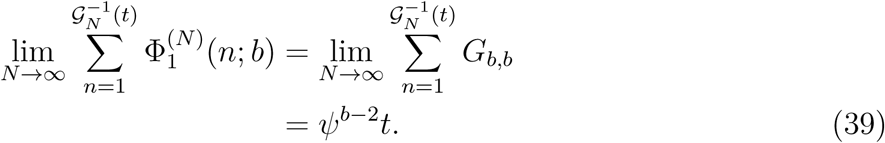

Hence, Theorem 1 implies that for each sample size *k ∈* ℕ the limit of the time-scaled ancestral process 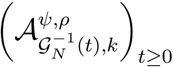 exists and from Equation 39 it follows that 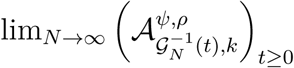 is a time-homogeneous Λ *- k-*coalescent. Further,

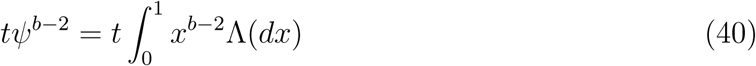

holds for all *b ∈* ℕ if and only if Λ is the Dirac measure at *ψ*. Thus, in the large population-size limit the ancestral process, 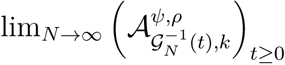, converges to a psi-*k*-coalescent - 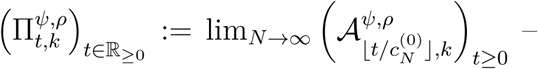 that is equal (in distribution) to a regular psi-*k*-converges 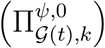 with time (non-linearly) rescaled by

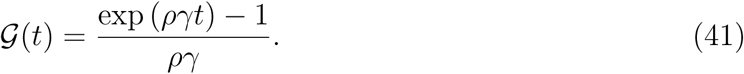

Put differently, analogous to the results obtained for the Kingman coalescent (Griffiths and Tavaré 1994; Griffiths and Tavaré 1998; Kaj and Krone 2003), the time-inhomogeneous ancestral limiting process of the extended Moran model with exponential growth can be transformed into a time-homogeneous psi-coalescent with coalescent rates given by Equation 10 with branches rescaled by Equation 41 allowing it to be simulated easily and efficiently. Intuitively, the transformation sums over the coalescence intensities of the time-inhomogeneous process and weighs them by the time they were effective, such that on the new time-scale coalescent intensities are constant across time, and the (re-scaled) process is time-homogeneous (see also Kaj and Krone 2003). Thus, changing the time-scale by Equation 41 compensates for the shrinking population sizes (going backwards in time) and the effect of increasing (total) coalescent rates.

To highlight the duality between the two processes – i.e., the (forward in time) extended Moran model and the corresponding coalescent – key properties (e.g., the summed length of all branches with *i* descendants *T*_*i*_ and the total tree length *T*_tot_) are compared in the Supporting Information (**Extended Moran model simulations**). Finally note, that Equation 41 is – except for the additional factor *γ* that is proportional to the coalescent time scale – structurally identical to the time-change function in the Kingman case (Griffiths and Tavaré 1998). However, since 𝒢 (*t*) depends on the product 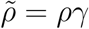, it is impossible to obtain a direct estimate of *ρ* (or *γ*) without additional information, and thus – analogous to the case of the population scaled mutation rate *θ* – only the compound parameter 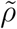 can be estimated. To keep notation simple though, we will refer to *ρ* (instead of 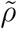) when referring to growth rate estimates.

### Joint inference of coalescent parameters and population growth rates

In this paragraph we modify the likelihood function

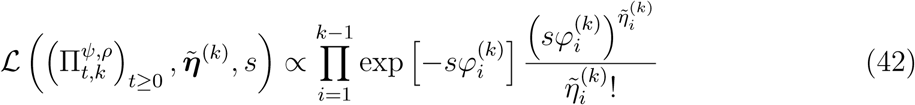

derived in the Model and Methods section to jointly infer the coalescent parameter *ψ* and the population growth rate *ρ*. Note that while the general form of the likelihood function (eq. 14) is independent of the generating coalescent process, changes in *ψ* and *ρ* affect the normalized expected SFS as given by

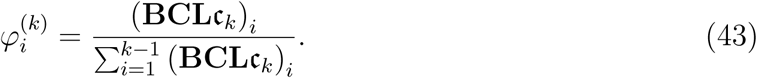

Recall that **B** and **C** depend neither on *ψ* nor *ρ*, and that **L** does depend on *ψ* but not on *ρ*, and that the time-inhomogeneity of the underlying coalescent process only enters through the first coalescence times c_*k*_, which are given by

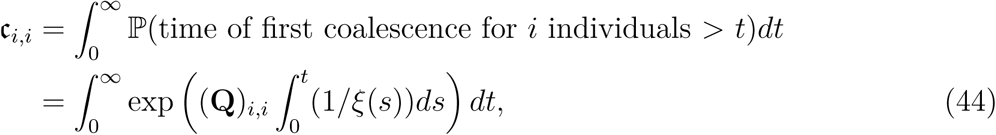

where *ξ*(*s*) denotes the intensity measure (Polanski and Kimmel 2003; Bhaskar *et al.* 2015; Spence *et al.* 2016). For the psi-coalescent with exponential growth *ξ*(*t*) = *e*^*−ργt*^ such that Equation 44 becomes

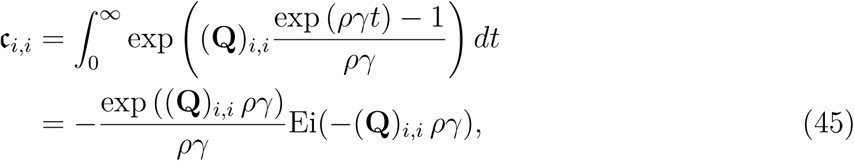

where 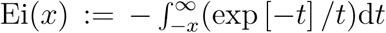 denotes the exponential integral. Thus, when growth rates are measured on their corresponding coalescent scale – i.e., *ργ* under the psi-coalescent versus *ρ* under the Kingman coalescent – Equation 45 is a generalization of the Kingman coalescent result (eq. 17) derived by Polanski and Kimmel (2003). Finally, combining Equation 45 with Equation 43 allows for the exact computation of the normalized expected SFS φ^(*k*)^, avoiding the simulation error that would be introduced by Monte Carlo simulations.

Figure 2 shows the normalized expected SFS obtained from Equation 43, where higher frequency classes have been aggregated (i.e., lumped) for different values of *ψ* and *ρ*. In line with previous findings, both multiple mergers and population growth lead to an excess in singletons (Durrett and Schweinsberg 2005; Eldon *et al.* 2015; Niwa *et al.* 2016). Furthermore, this excess increases as sample size increases under the psi-coalescent (Fig. SI C_1), while it decreases for the Kingman coalescent independent of the presence or absence of exponential growth. These qualitative differences stem from the different footprints reproductive skew and exponential growth leave on a genealogy. While the latter is a simple rescaling of branch lengths leaving the topology unchanged, multiple-merger coalescents by definition affect the topology of the genealogical tree (Eldon *et al.* 2015). In particular, when *ψ* is large, adding samples will disproportionally increase the number of external branches *T*_1_ such that the genealogy will become more star-like, rendering disproportionately more singletons.

**Figure 2.**
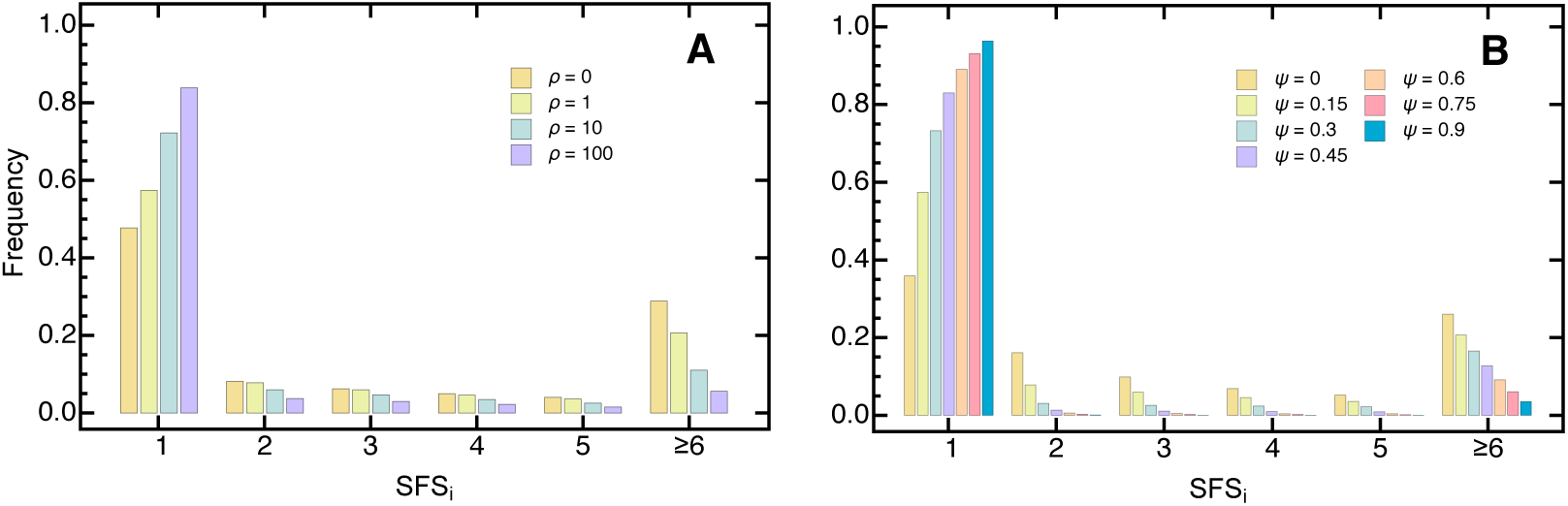
The normalized expected (lumped) SFS for the psi-coalescent for an exponentially growing population (eq. 18) with sample size *k* = 20 (**A**) for different values of *ρ* and fixed *ψ* = 0.15 and (**B**) for different values of *ψ* and fixed *ρ* = 1. The sixth entry in the SFS contains the aggregate of the higher frequency classes.

Though the excess in singletons characterizes either process, their higher frequency classes will typically differ (Eldon *et al.* 2015). When both processes – reproductive skew and exponential growth – act simultaneously though, their joint effects on the SFS (non-trivially) combine. As expected, increasing growth under the psi-coalescent further exacerbates the excess in singletons. More generally, exponential growth leads to a systematic left shift in the SFS towards lower frequency classes that is independent of *ψ*. Increasing *ψ* on the other hand changes the SFS – and in particular the higher frequency classes – non-monotonically even if there is no population growth (Fig. SI C_2). Interestingly, for *ρ* = 0 the last entry of the normalized expected SFS 𝔼 [*η*_*k–*1_] initially increases with *ψ* and takes an intermediate maximum, decreases monotonically until *ψ ≈* 0.85, peaks again and then quickly reduces to 0 as *ψ* approaches 1. This effect prevails as sample size increases (Fig. SI C_2) even though the intermediate maximum slightly shifts towards lower *ψ*. However, this intermediate maximum is effectively washed out by increasing *ρ* such that the second peak becomes the maximum. Furthermore, the shape of the peak becomes more pronounced as sample size increases. Thus, reproductive skew and exponential growth leave complex and distinct genomic footprints on the SFS. While in theory population growth and reproductive skew should be identifiable, this in practice strongly depends on sample size (Spence *et al.* 2016). In the next section we will assess the accuracy of our joint estimation framework and perform extensive validation (eq. 14) on large-scale simulated data.

### Simulated coalescent and demographic models

To test our inference framework, we followed two different simulation approaches, each corresponding to two biological limiting cases. In both, data was simulated for the Cartesian product set over *ψ* = {0, 0.15, 0.3, 0.45, 0.6, 0.75, 0.9}, *ρ* = {0, 1, 10, 100}, *k* ={20, 50, 100, 200} and *s* ={100, 1 000, 10 000} per locus over 10,000 replicates each. In order to make results comparable across different coalescent models and thus across different values of *ψ* and *ρ*, we calculated the population-scaled mutation rate *θ* based on Watterson’s estimator (Watterson 1975),

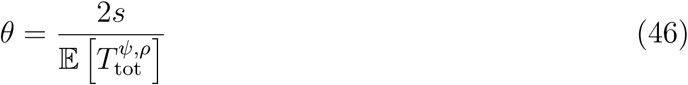

for a fixed number of segregating sites *s* over the expected total tree length under the generating coalescent model (given by the denominator in eq. 43). Note that *T*_tot_ decreases with both increasing *ψ* and *ρ*. Thus, keeping *s* constant implies that *θ* effectively increases with ψ and *ρ*. We will discuss the latter point in more detail in light of the results below. Data was simulated for the following two underlying genetic architectures:

- Case 1 (Independent-sites simulations): Under the Poisson random field assumption, the underlying coalescent tree at each site is independent (Sawyer and Hartl 1992; Bhaskar *et al.* 2015). Thus, by averaging over independent realizations of the (shared) underlying coalescent process the SFS can be obtained by randomly drawing from a multinomial distribution such that ***η*** *∼* Multinomial (*s, φ*).
- Case 2 (Whole-genome simulations): In this scenario we consider a genome of *l* = 100 and *l* = 1,000 independent loci, respectively, where sites within each locus share the same genealogy (i.e., coalescent tree). Thus, for each locus we draw a random genealogy according to equations 10 and 41, superimpose *s ∼* Poisson(*θ/*2) random mutations onto the ancestral tree by multinomial sampling, and aggregate the individual locus SFS into a single genome-wide SFS.

Finally, data sets where *s* = *η*_1_ (i.e., where all segregating sites were singletons) were discarded and simulated again since these do not allow the underlying coalescent parameter and demographic history to be identified.

### Accuracy of joint estimation framework

Next we evaluated the accuracy of the joint estimation framework by means of the mean absolute deviation (MAD) 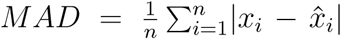, the mean deviation 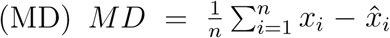, the mean squared error (MSE) 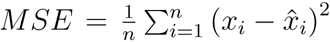, and the median deviation (MDD) where *x* and 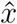 denote the true and the estimated parameter, respectively. If not stated otherwise, results in the main text are shown for the default parameters *k* = 100 and *θ* (eq. 46) with *s* = 10,000. More results are given in Supporting Information (**Support**ing Figures) and (**Supporting Tables**).

#### Inference under the independent-sites assumption

First, for a consistency check we applied our grid-search algorithm to estimate *ψ* and *ρ* from an idealized SFS (i.e., where the SFS accurately reflects the expected branch length under the generating coalescent and demographic model ***φ*** except for distortions due to rounding). An exemplary likelihood surface (eq. 14) for such an idealized SFS is depicted in Figure 3, which shows that the likelihood surface – up to the resolution of the grid point – is smooth and generally unimodal and that the true parameters can be estimated accurately. Furthermore, Figure 3 shows that there is generally a negative correlation between *ψ* and *ρ*, and that the likelihood surface tends to be steeper and more concentrated along the *ψ* direction, which suggests that growth rate estimates might show a larger variance and could in general be more difficult to estimate. The steepness of the likelihood surface along the *ψ* axis tends to increase with *ψ* and sample size *k*, suggesting that the accuracy for estimating *ψ* should increase as well, while it should become more difficult to estimate *ρ* accurately.

**Figure 3.**
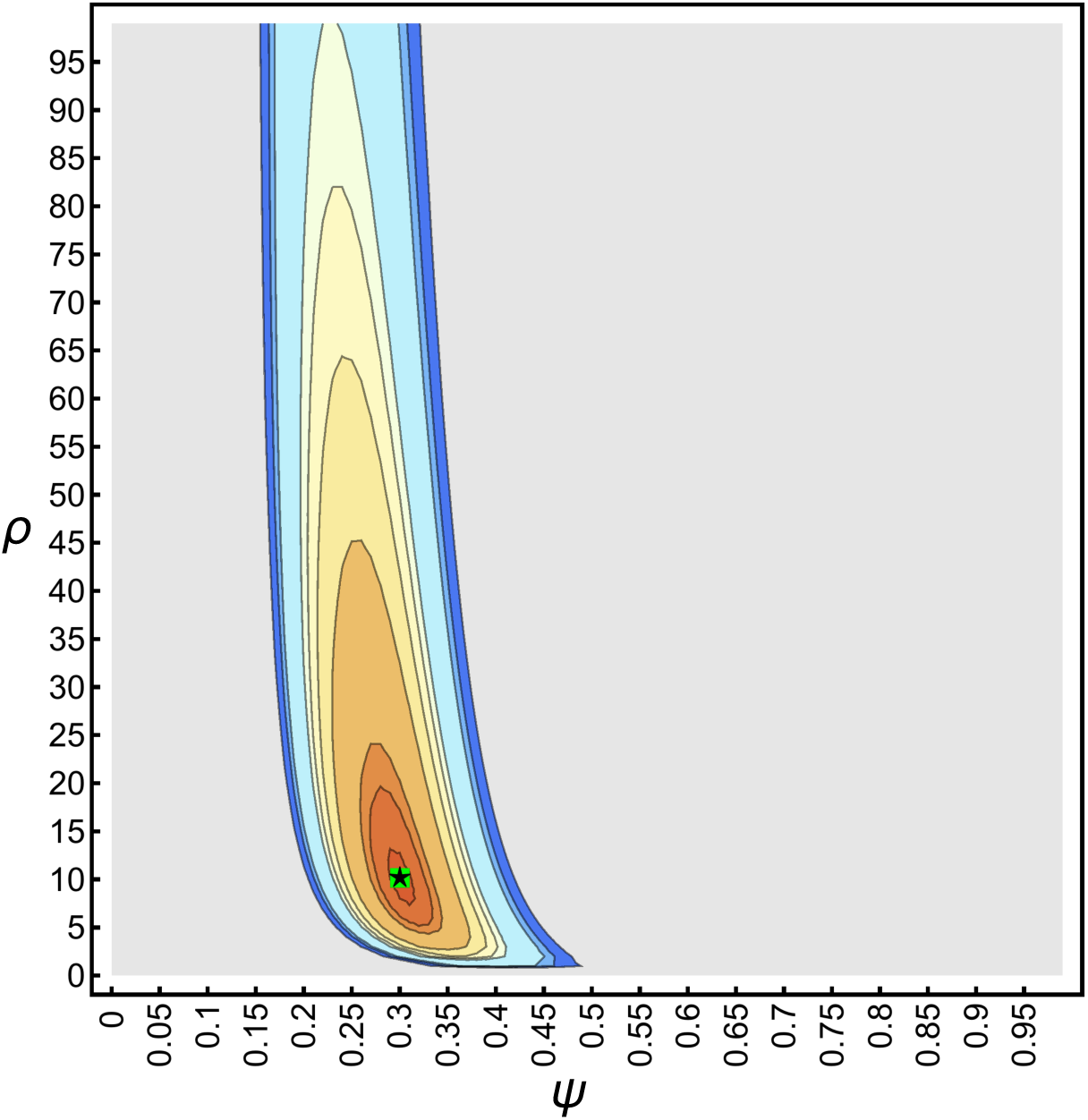
Likelihood surface (eq. 14) of the idealized SFS with *k* = 100, *ψ* = 0.3, *ρ* = 10 and *s* = 10,000. Contours show the 0.95, 0.9675, 0.975, 0.99, 0.99225, 0.9945, 0.99675, 0.999, 0.99945 and 0.9999 quantiles. Likelihoods below the 0.95 quantile are uniformly colored in gray. The green square shows the true *ψ* and *ρ*. The black star shows the maximum likelihood estimates 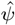 and 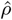.

An exemplary distribution of the jointly inferred maximum likelihood estimates 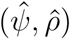 assuming independent sites is shown in Figure 4. The shape of this distribution resembles that of the likelihood surface (Fig. 3), indicating that there is some variance – in particular along the *ρ*-axis – in the maximum likelihood estimates. However, the median and the mean of the distribution match the true underlying coalescent and growth rate parameters (i.e., *ψ* and *ρ*) very well, implying that if sites are independent, 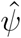 and 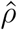 are unbiased estimators.

**Figure 4.**
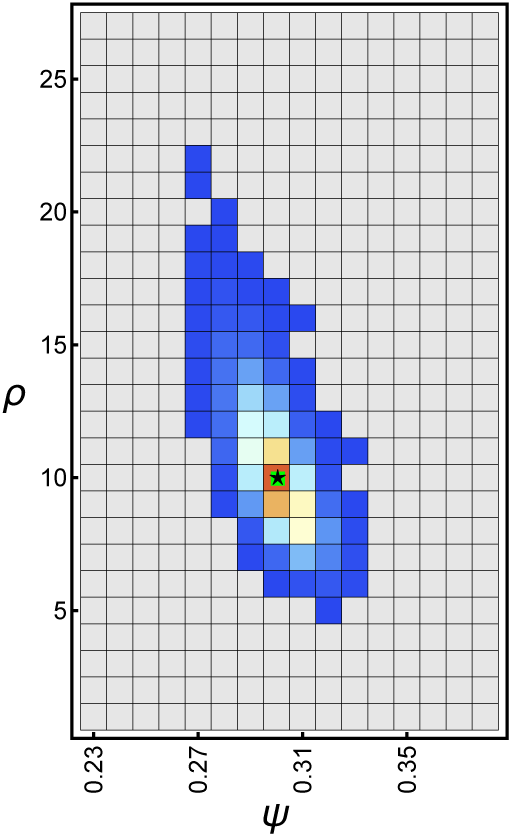
Heatplot of the frequency of the maximum likelihood estimates for 10,000 data sets assuming independent sites with *k* = 100, *ψ* = 0.3, *ρ* = 10, and *θ* (eq. 46) with *s* = 10,000. Counts increase from blue to red with grey squares showing zero counts. The green square shows the true *ψ* and *ρ*. The black star shows the median (and mean) of the maximum likelihood estimates 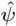 and 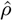.

Generally, as expected from the shape of the likelihood surface, *ψ* is estimated with high accuracy and precision, even for large sample sizes (*k* = 200) with only a few segregating sites (*s* = 100) and (nearly) independent of *ρ* (Fig. 5A, SI C_3A, SI C_4A; Table SI D_1). Growth rate estimates 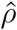, however, show a larger variance and for some parameters – namely large *k* and small *s* – might be slightly upwardly biased when both the coalescent parameter and the growth rate are large (Fig. 5B, SI C_3B, SI C_4B; Table SI D_2). Though, as the number of segregating sites increases, this bias vanishes and the variance decreases (Fig. SI C_5), highlighting that the joint estimation procedure gives asymptotically unbiased estimators.

**Figure 5.**
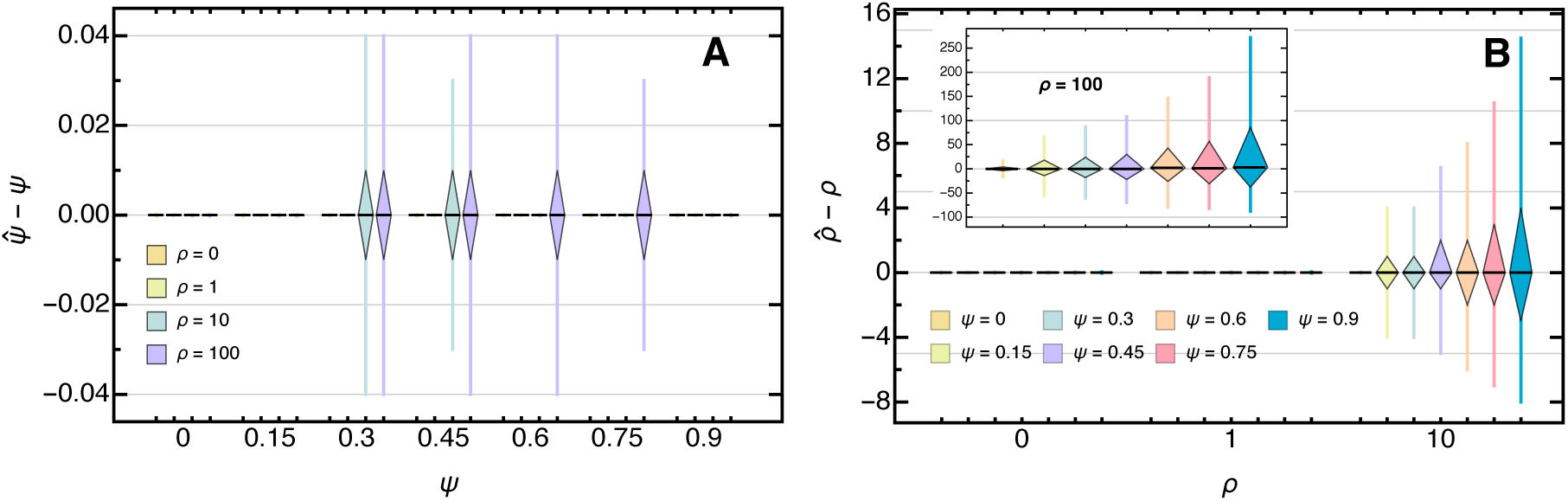
Boxplot of the deviation of the maximum likelihood estimate from the true (**A**) *ψ* and (**B**) *ρ* for 10,000 data sets assuming independent site with *k* = 100 and *θ* (eq. 46) with *s* = 10,000. Boxes represent the interquartile range (i.e., the 50% C.I.) and whiskers extend to the highest/lowest data point within the box *±*1.5 times the interquartile range.

For a given *s*, increasing sample size *k* increases the signal-to-noise ratio and thus the error in both 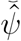 and 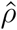 (Tables SI D_1, SI D_2, SI D_3, SI D_3) which is most noticeable in growth rate estimates, in particular when *ρ* is large (Fig. SI C_6). This increase in estimation error can (partially) be compensated by increasing the number of segregating sites *s* (Fig. SI C_7, Table SI D_5). Specifically, if the true underlying *ψ* is large (i.e., if the offspring distribution is heavily skewed) an increasing number of segregating sites is needed to accurately infer *ρ*. However, the total tree length 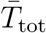 – and thus the number of segregating sites *s* – is expected to decrease sharply with *ψ* (Eldon and Wakeley 2006), implying that trees tend to become shorter under heavily skewed offspring distributions. This effect could (again, partially) be overcome by increasing sample size since 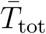 – unlike the Kingman coalescent – scales linearly with *k* as *ψ* approaches 1 (Eldon and Wakeley 2006). However, population growth will reduce 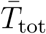 and the number of segregating sites even further.

Calculating *θ* based on a fixed and constant (expected) number of segregating sites for the assessment of the accuracy of the estimation method evades this problem to some extent. However, by making this assumption we effectively increase *θ* in our simulations as *ψ* and *ρ* increases. Our results suggest, though, that even more segregating sites than considered in this study (i.e., an even larger *θ*) would be necessary to infer population growth accurately. Thus, unless (effective) population sizes and/or genome-wide mutation rates are large it might be very difficult to infer population growth if the offspring distribution is heavily skewed (i.e.,. if *ψ* is large). On the other hand, the few studies that have estimated *ψ* generally found it to be small (Eldon and Wakeley 2006; Birkner *et al.* 2013; Árnason and Halldórsdóttir 2015) leaving it unresolved whether this problem is of any practical importance when studying natural populations.

#### Inference from genome-wide data

We next tested the accuracy of our joint estimation framework when applied to genome-wide data obtained from *l* = 100 independent loci. An exemplary distribution of the jointly inferred maximum likelihood estimates 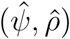 is depicted in Figure 6; Figure 7 shows the overall performance of the joint estimation method when applied to genome-wide data. While the whole-genome simulations are designed such that each site in a given locus shares the same underlying genealogy, and thus violate the assumption of (statistical) independence between sites, we find that coalescent and growth rate parameters (i.e., *ψ* and *ρ*) can be estimated robustly and accurately. In concordance with the independent-sites simulations, the variance in 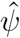 is typically small, whereas 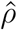 spreads considerably, and increasingly so if *ψ* is large. The mean and the median of the coalescent parameter and growth rate estimates are again centered around the true value, implying that 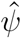 and 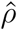 are unbiased estimators (see also Tables SI D_11, SI D_12, SI D_13, SI D_14).

**Figure 6.**
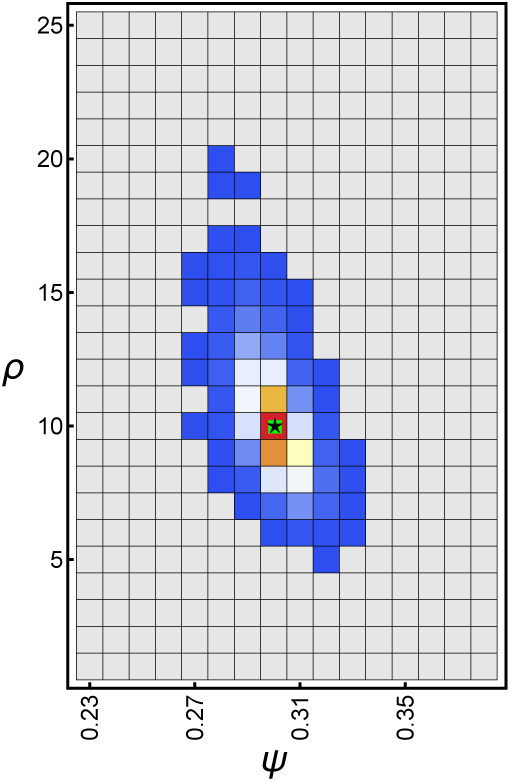
Heatplot of the frequency of the maximum likelihood estimates for 10,000 whole-genome data sets assuming with *l* = 100*, k* = 100, *ψ* = 0.3, *ρ* = 10*, γ* = 1.5 and *θ* (eq. 46) with *s* = 1,000. Counts increase from blue to red with grey squares showing zero counts. The green square shows the true *ψ* and *ρ*. The black star shows the median (and mean) of the maximum likelihood estimates 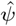 and 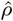.

**Figure 7.**
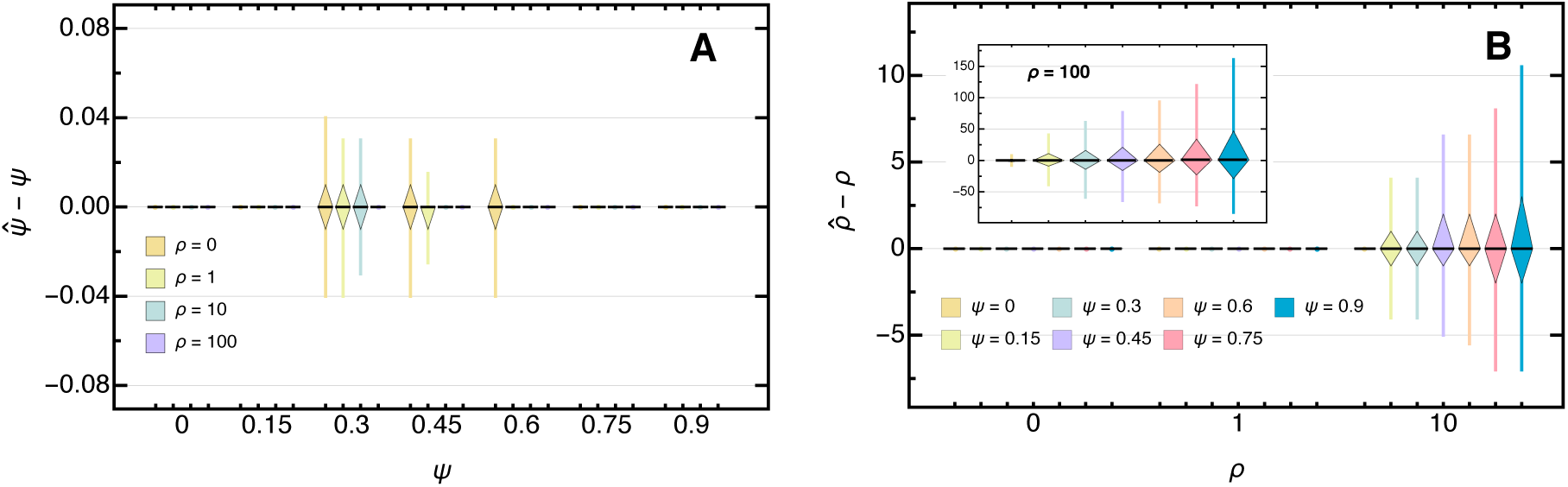
Boxplot of the deviation of the maximum likelihood estimate from the true (**A**) *ψ* and (**B**) *ρ* for 10,000 whole-genome data sets with *l* = 100*, k* = 100, *γ* = 1.5, and *θ* (eq. 46) with *s* = 1,000. Boxes represent the interquartile range (i.e., the 50% C.I.) and whiskers extend to the highest/lowest data point within the box *±*1.5 times the interquartile range.

Interestingly, the precision of the coalescent and growth rate parameter estimates increases along with the number of loci (i.e., the number of independent coalescent realizations) while keeping the number of segregating sites constant (Table SI D_10). While this effect is to some extent expected as increasing the number of (independent) loci reduces the approximation error (see above), it suggests that sequencing efforts should be put on covering the genome in its entirety rather than on increasing coverage of individual genomic regions.

#### Distance-based inference and the effect of lumping

As an alternative to the likelihood-based method, Eldon *et al.* (2015) proposed an ABC approach based on a minimum-distance statistic (eq. 15). In this section we assess the accuracy of 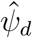 and 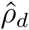 when estimated from *d*_1_ and *d*_2_ distances (i.e., the *l*_1_ and *l*_2_ distance). A surface plot of the *l*_1_ and the *l*_2_ distance is shown in Figure SI C_8. We find that for the *l*_1_ and the *l*_2_ distance results are comparable to those of the likelihood-based estimates, but generally display a larger variance (Fig. SI C_9, SI C_10, SI C_11). Likelihood-based estimates 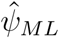 tend to be more accurate across the entire parameter space, though differences between the two are marginal.

Over the majority of the parameter space the same holds true for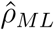. Particularly for small to intermediate *ψ* the likelihood-based approach outperforms both distance-based approaches considerably (Table SI D_7). Interestingly though, for large *ψ* and *ρ* (i.e., in the part of the parameter space where estimating *ρ* is generally difficult) the *l*_1_ distance approach gives more accurate estimates. When increasing the number of segregating sites, though, the likelihood approach becomes more accurate again, suggesting that the *l*_1_ distance-based approach only outperforms the likelihood-based approach when there is insufficient data (not shown). These general findings are also upheld when considering genome-wide data (Figs. SI C_12, SI C_13). Despite the slightly reduced power as compared to the maximum likelihood approach, our results indicate that, given the asymptotic properties, both the *l*_1_ and the *l*_2_ distance should perform reasonably well when used in a rejection-based ABC analysis.

Finally, we investigated the effect of lumping (i.e., aggregating the higher-frequency classes of the SFS into a single entry after a given threshold *i*) on the performance of our estimator. In contrast to Eldon *et al.* (2015), who found that lumping can improve the power to distinguish between multiple-merger coalescent models and models of population growth, we find that estimates based on the lumped SFS (using *i* = 5 and *i* = 15) show considerably more error (Tables SI D_8, SI D_9). While *ψ* can again be reasonably well estimated, 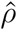 – in particular when *ψ* and/or *ρ* are large – is orders of magnitude more inaccurate when higher frequency classes are lumped. The reason is that when trying to differentiate between different coalescent or growth models, lumping can reduce the noise associated with the individual higher frequency classes and thus increases the power, provided that the different candidate models show different mean behaviors in the lumped classes (Eldon *et al.* 2015). While this seems to hold true when considering “pure” coalescent or growth models, the joint footprints of skewed offspring distributions and (exponential) population growth are more subtle. In particular, since growth induces a systematic left shift in the SFS towards lower frequency classes, most of the information to distinguish between a psi-coalescent with or without growth is lost when aggregated.

### Mis-inference of coalescent parameters when neglecting demography

As argued above, both reproductive skew and population growth result in an excess of singletons (i.e., low-frequency mutations) in the SFS. However, topological differences between the two generating processes in the right tail of the SFS allows distinguishing between the two. In particular, fitting an exponential growth model and not accounting for reproductive skewness results in a vastly (and often unrealistically) overestimated growth rate (Eldon *et al.* 2015).

Here, we investigate how coalescent parameter estimates (i.e., 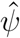) are affected when not accounting for (exponential) population growth (i.e., assuming *ρ* = 0) when both processes act simultaneously. As expected we find that 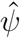 is consistently overestimated (Fig. 8) and that the estimation error – independent of *ψ* – increases with larger (unaccounted for) growth rates. This is because, unless the underlying genealogy is star-shaped (e.g., when *ψ* = 1), growth will always left-shift the SFS and hence increase the singleton class. Thus, when assuming *ρ* = 0, increasing *ψ* compensates for the “missing” singletons. Interestingly though, the estimation error changes non-monotonically with *ψ*, and for large *ρ* can be as great as twice the value of the true underlying coalescent parameter. Furthermore, for low to intermediate *Ψ*, even small growth rates can result in a relative error of up to 23%. Overall, not accounting for demography can lead to serious biases in *ψ* with broad ecological implications when trying to understand the variation in reproductive success.

**Figure 8.**
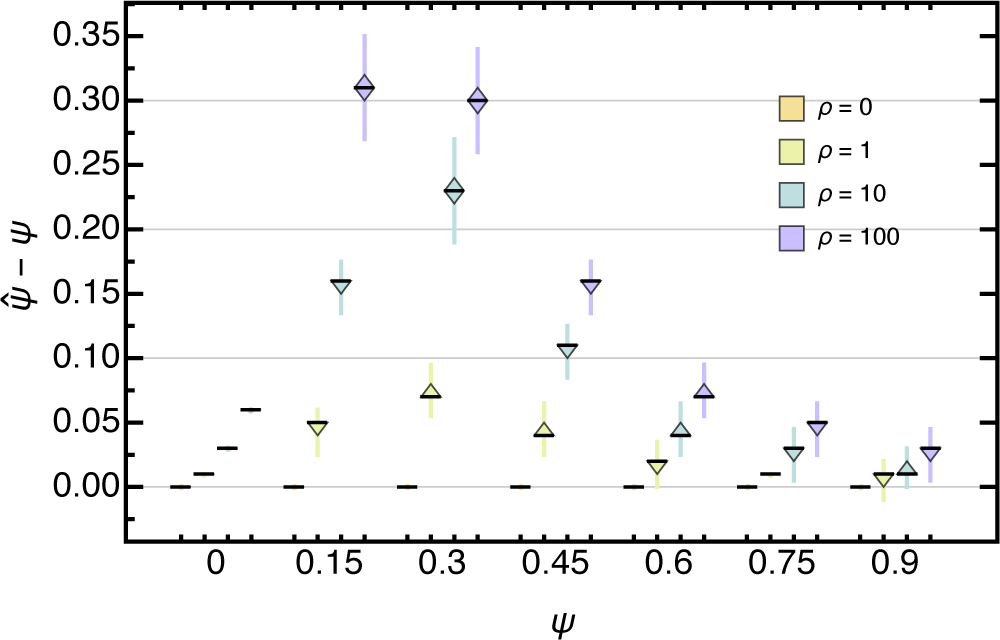
Boxplot of the deviation of the maximum likelihood estimate from the true *ψ* for 10,000 data sets assuming independent sites with *k* = 100 and *θ* (eq. 46) with *s* = 10,000. when not accounting for population growth. Boxes represent the interquartile range (i.e., the 50% C.I.) and whiskers extend to the highest/lowest data point within the box *±*1.5 times the interquartile range.

### Application to sardine data

Finally, we applied our joint inference framework to a derived SFS for the control region of mtDNA in Japanese sardine (*Sardinops melanostictus*; File E1). Niwa *et al.* (2016) recently analyzed this data to test whether the observed excess in singletons was more likely caused by a recent population expansion or by sweepstake reproductive events and found that the latter is the more likely explanation. However, there is of course no *a priori* reason to believe that both reproductive skew and population growth could not have acted simultaneously.

When estimated jointly, the maximum likelihood estimate is 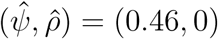, which implies considerable reproductive skew, but no (exponential) population growth (Fig. 9; see FigSI. SI C_14 for the corresponding *l*_1_ and *l*_2_ distance estimates). While our analysis confirms their results at first glance there are two points that warrant caution with this interpretation. First, as indicated by the contour lines in the plot there is some probability that the Japanese sardine population underwent a recent population expansion, though if it did, it only grew at a very low rate. Second, our inference is based on a single non-recombining locus (i.e., mtDNA) implying that there is correlation between sites. Our approximation, though, is exact only if there is independence between sites. While violations of the independence assumption seem to be robust on the genome-wide scale (see above; Fig. 7), per-locus estimates can vary drastically and might not be representative for the true underlying coalescent process (not shown).

**Figure 9.**
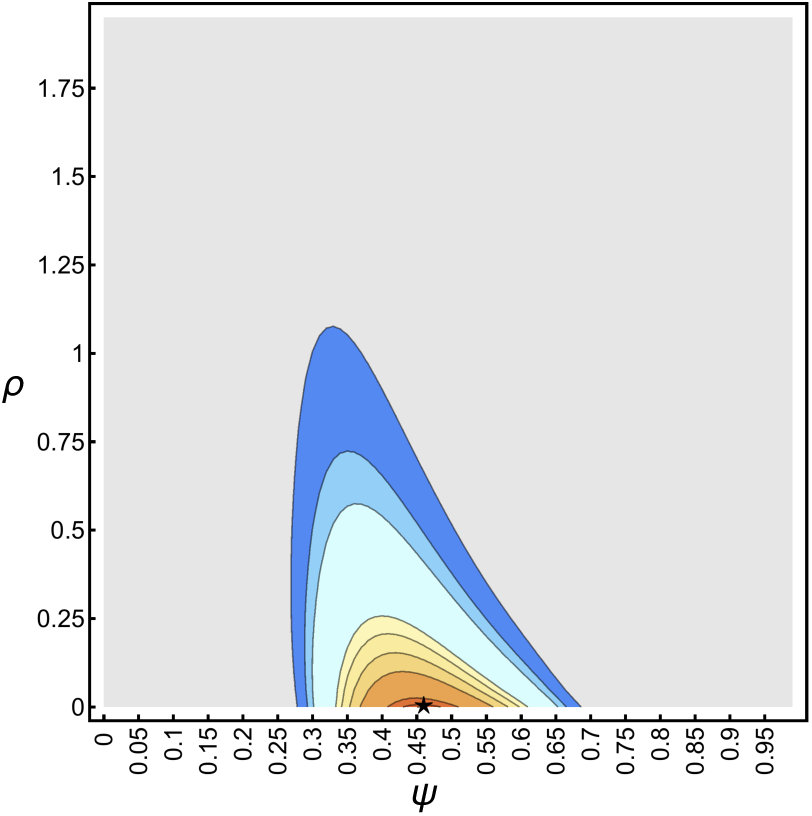
Likelihood surface (eq. 14) of the unfolded SFS (given the ML rooted tree) of the sardine mtDNA sequences with *k* = 106 and *s* = 78. Contours show the 0.95, 0.9675, 0.975, 0.99, 0.99225, 0.9945, 0.99675, 0.999, 0.99945 and 0.9999 quantiles. Likelihoods below the 0.95 quantile are uniformly colored in gray. The black star shows the maximum likelihood estimates 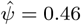 and 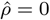.

### Concluding remarks

This study marks the first multiple-merger coalescent with time-varying population sizes derived from a discrete time random mating model, and provides the first in-depth analyses of the joint inference of coalescent and demographic parameters. Since the Kingman coalescent represents a special case of the general class of multiple-merger coalescents (Sagitov 1999; Pitman 1999; Schweinsberg 2000; Donnelly and Kurtz 1999; Spence *et al.* 2016), it is interesting and encouraging to see that our analytical results – i.e., the time-change function (eq. 41) and the first expected coalescence times (eq. 45) – are generalizations of results derived for the Kingman coalescent (Griffiths and Tavaré 1998; Polanski and Kimmel 2003). In fact, when growth rates are measured within the corresponding coalescent framework (e.g., as 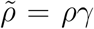 for the psi-coalescent) these formulas should extend to other, more general multiple-merger coalescents. This also holds true for the challenges arising when calculating the normalized expected SFS (eq. 13) which is central to estimating coalescent parameters and growth rates: Because of catastrophic cancellation errors – mainly due to summing over alternating sums and numerical representations of the exponential integral Ei(*x*) – computations have to be carried out using multi-precision libraries (Spence *et al.* 2016).

While both *ψ* and *ρ* can generally be estimated precisely, accurate estimation of the latter requires sufficient information (i.e., a large number of segregating sites) especially when offspring distributions are heavily skewed (i.e., if *ψ* is large). However, since strong recurrent sweepstake reproductive events – analogous to recurrent selective sweeps – constantly erase genetic variation (i.e., reduce the number of segregating sites), there might be little power to accurately infer *ρ* in natural populations in these cases. In accordance with previous findings derived for the Kingman coalescent (Terhorst and Song 2015), increasing sample size does not improve the accuracy of demographic inference (i.e., estimating *ρ*) for a fixed (expected) number of segregating sites *s*. However, unlike in the Kingman coalescent where *s* increases logarithmically with sample size, genetic variation in *ψ* increases linearly for large *ψ* which could offset – or at least the hamper – this effect.

More importantly, these results have proven to be robust to violations of the assumptions underlying the approximate likelihood framework (eq. 14), namely that the expectation of a ratio can be approximated by the ratio of two expectations (i.e., 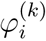), allowing *ψ* and *ρ* to be estimated accurately on a genome-wide scale. Interestingly, the performance of the estimators seemed to improve when considering more independent loci (while keeping the number of segregating sites constant). Note though, that we have used a very simplistic genetic architecture, in particular one where sites within each locus are maximally dependent and there is no correlation among genealogies across different loci (i.e., where loci are independent). While these assumptions might be met for some loci and sites, they generally mark the endpoint of a continuum of correlations. Importantly, these linkage (dis)equilibria (i.e., the extent of statistical independence between sites) depend not only on the rate of recombination but also on the specifics of the reproduction parameters – and can potentially be elevated despite frequent recombination or largely absent despite infrequent recombination in the MMC setting (Eldon and Wakeley 2008; Birkner *et al.* 2012), potentially biasing results. For instance, when trying to estimate the duration and the rate of exponential growth under the Kingman coalescent, Bhaskar *et al.* (2015) found that linkage equilibria cause the approximate likelihood equation(eq. 14) to become increasingly inaccurate, and thus bias estimates. Likewise, Schrider *et al.* (2016) recently found that linked positive selection can severely bias demographic estimates. While their analyses assumed a Kingman framework, positive selection and recurrent selective sweeps typically result in multiple merger events (Durrett and Schweinsberg 2004, 2005; Neher and Hallatschek 2013). Thus, if neutral regions are tightly linked to selected site they will – at least partially – share the genealogical relationship with the selected region and potentially skew inference. Despite the fact that our model here considers organisms with skewed offspring distributions under neutrality owing to the specifics of their reproductive biology, increasing *ψ* is tantamount to increasing the strength of positive selection under a non-neutral model, which is thus relevant to a very broad class of organisms indeed. It is important to note that while both processes – selection and sweepstake reproductive events – have a similar effect on the SFS (i.e., an excess of low-frequency alleles and a slight increase in high-frequency alleles), there are of course vast qualitative differences in the underlying processes and their causes. First, in the presence of selection, offspring no longer choose their parents at random, such that selected alleles need to be tracked along the genealogy (e.g., see the ancestral selection graph under the Kingman coalescent (Krone and Neuhauser 1997) or under the Λ-coalescent (Etheridge *et al.* 2010)). Second, similar to the effects of demography, sweepstake reproductive events should have a genome-wide impact, whereas traces of selection should remain local, unless selection is very strong such that only a single individual gives rise to the entire next generation. Thus, it should in principle be possible to discriminate between the two processes, though, also analogous to demography, it will be important to investigate the conditions under which positively selected loci will be expected to reside in the tails of genomic distributions under such models (see Thornton and Jensen 2007).

Overall, our analyses emphasize the importance of accounting for demography and illuminates the serious biases that can arise in the inferred coalescent model if ignored. Such bias can have broad implications on inferred patterns of genetic variation (Eldon and Wakeley 2006; Tellier and Lemaire 2014; Niwa *et al.* 2016), including misguiding conservation efforts (Montano 2016), and obscuring the extent of reproductive skew.

Finally, most of the current analytical and computational tools have been derived and developed under the Kingman coalescent. In order to achieve the overall aim of generalizing the Kingman coalescent model (Wakeley 2013), these tools, though often computationally challenging, need to be extended. Great efforts have recently been undertaken towards developing a statistical inference framework allowing for model selection (Birkner and Blath 2008; Eldon 2011; Birkner *et al.* 2011, 2012, 2013; Steinrücken *et al.* 2013; Eldon *et al.* 2015; Spence *et al.* 2016). By setting up a discrete-time random mating model and deriving the ancestral process, along with providing the analytical tools necessary to enable the joint inference of offspring distribution and demography, this study makes an important contribution towards this goal.

## Acknowledgements

This project was funded by grants from the Swiss National Science Foundation (FNS) and a European Research Council (ERC) Starting Grant to JDJ. We thank Kristen Irwin, Adamandia Kapopoulou, Martin Möhle, Sylvain Mousset and Meike Wittmann for helpful discussion and comments on earlier versions of this manuscript, Hiro-Sato Niwa for providing the sardine data, and two anonymous reviewers for their insightful comments and constructive criticism which greatly helped to improve this manuscript.

## Supporting Information

### Detailed derivation of results

In this Supporting Information we will give a detailed derivation of the results which in the main text have been abbreviated to keep formulas concise. First, Equation 23 showing that 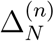 is bounded for all *n ∈* ℕ is derived as

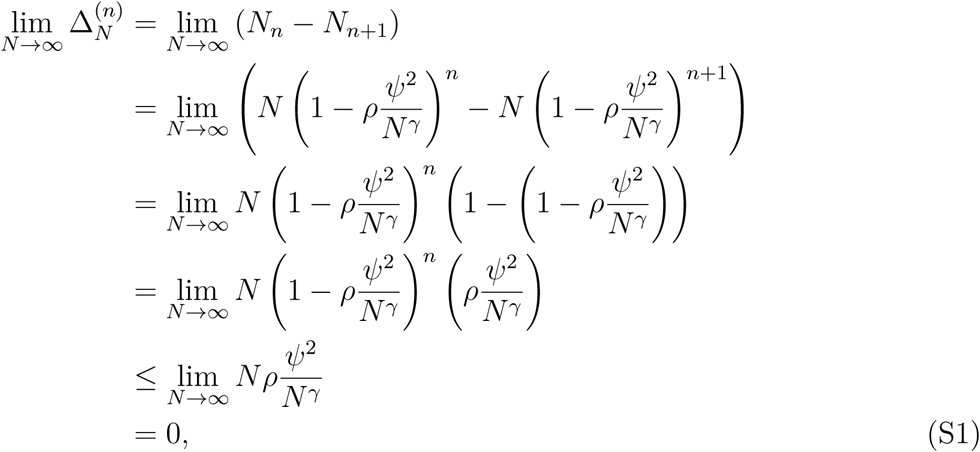

given *γ >* 1. Note that in the second to last line we have used that 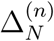 is always the largest for *n* = 0 (i.e., the population grows the largest from the previous to the current generation). Second, the step function *F*_*N*_ (*s*) (eq. 31) for the ancestral process is derived as

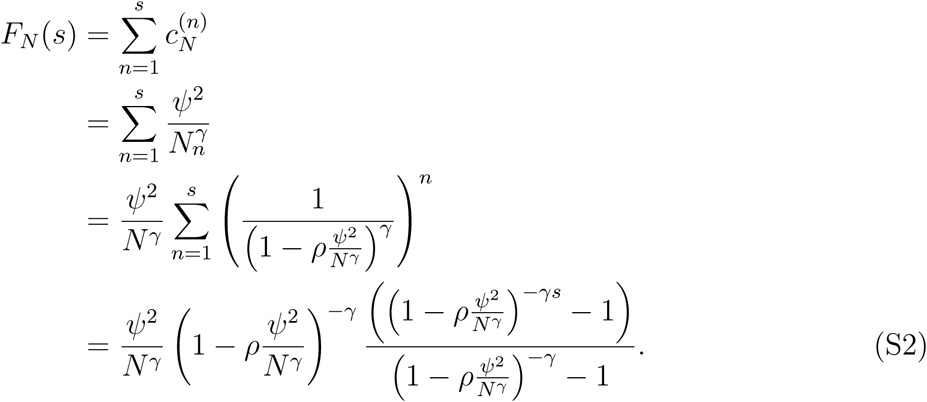

Then, by solving for *s* the time-change function 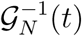 (eq. 32) for the ancestral process can be derived as

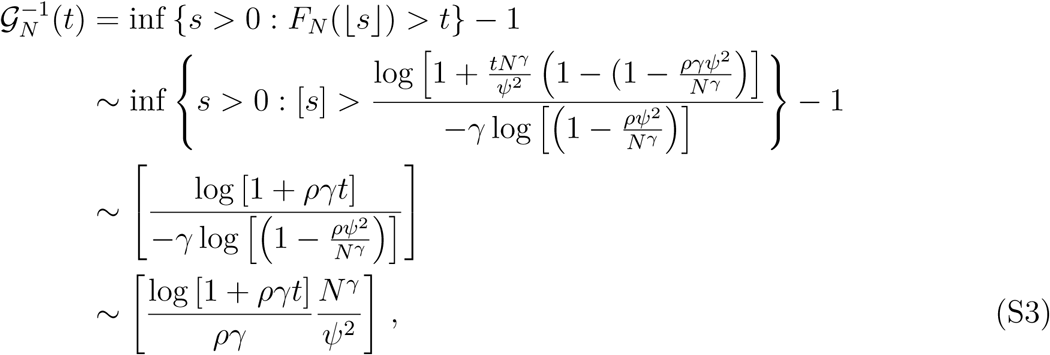

where we have used that 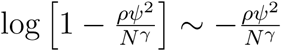 for sufficiently large *N*. Finally, the derivation showing that the ancestral process of the Moran model converges to a (time-inhomogeneous) psi-coalescent (eq. 39) is given by

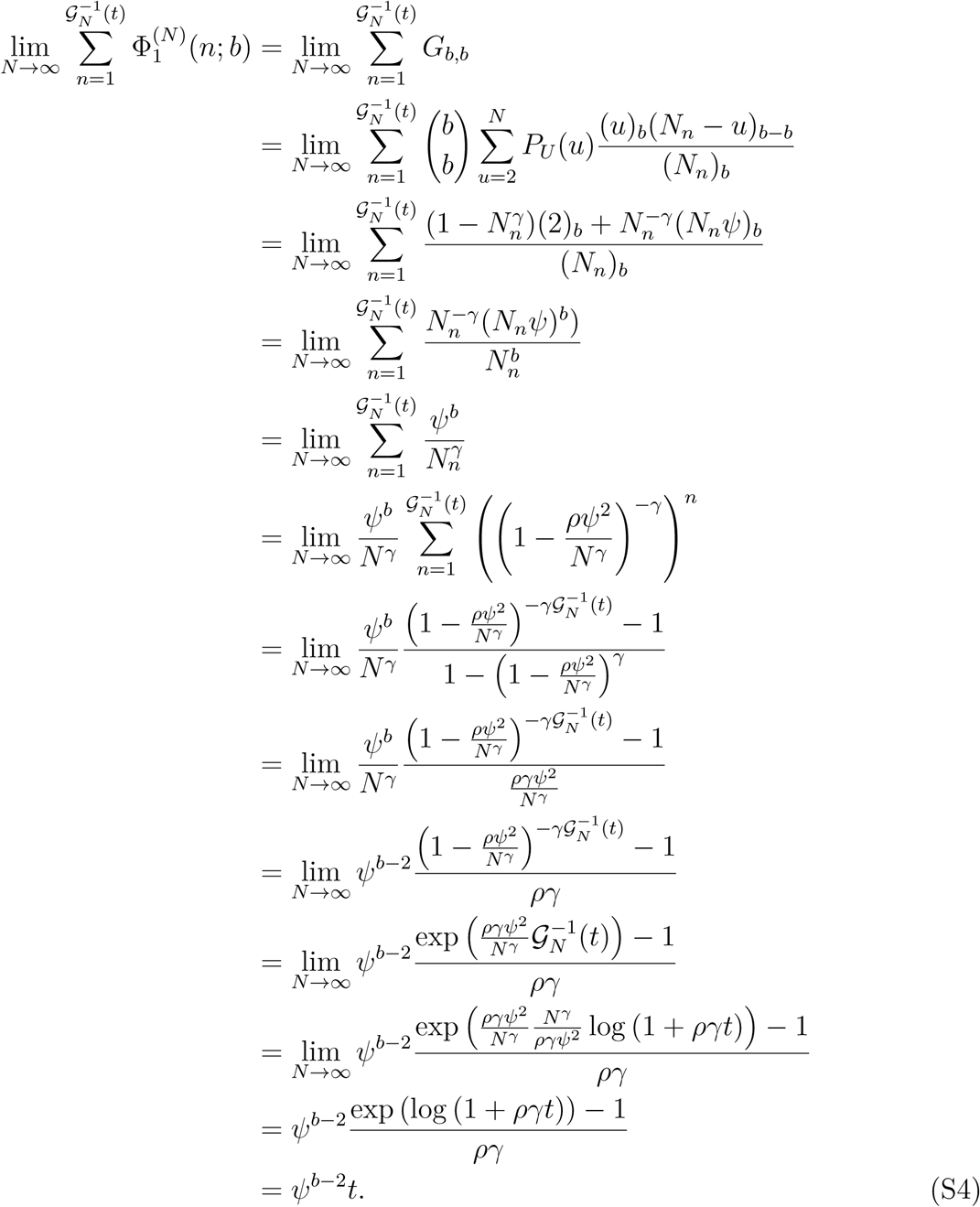

### Extended Moran model simulations

In this Supporting Information we compare a set of tree statistics obtained from the ancestral process 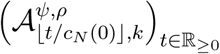 from the underlying extended Moran model to those obtained from the coalescent process 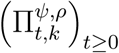 for different values of *ρ ∈* ℝ_*≥*0_, *ψ ∈* (0, 1) with *γ* = 1.5 (i.e., in the regime where sweepstake reproductive events dominate and the corresponding ancestral process is a time-inhomogeneous psi-coalescent). Coalescent simulations follow the algorithm outlined in the main text.

The rational behind the extended Moran model simulations is outlined in Figure SI B_1.

Starting from the present (i.e., *n* = 0) and choosing *k* random samples, these were followed successively backwards in time thus creating a coalescent tree from the leaves to the root. At each time step the population size was adjusted (shrinking backward in time; eq. 6) and the type of the reproductive event (eq. 5) along with the corresponding number of offspring *U*_*N*_ (*n*) (eq. 4) were (randomly) determined. Finally, to determine whether a merger event has occurred – i.e., if one of the *k* active lineages had found its parent in the previous time step – *U*_*N*_ (*n*) Bernoulli random numbers 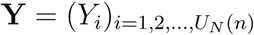 with parameter

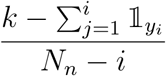

are drawn. Note that the denominator changes with the number of individuals that have not been assigned to a parent in the previous time step, while the nominator changes with the number of active lineages that have found their parent in the previous time step. In particular, 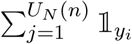 denotes the total number of active lineages that found their parent in the previous time step. Thus, if 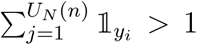 a coalescent event has occurred, and the number of active lineages in the next time step becomes 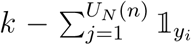. This process is repeated until *k* = 0, that is until all samples have found their most recent common ancestor. For both processes the total tree length *T*_tot_, the time to the most recent common ancestor *T M RCA*, and the ratio of the sum of the length of all branches with *i* descendants over the total tree length *T*_*i*_*/T*_tot_ were recorded. While the number of samples was limited to *k* = 4 for computational reasons, the match between coalescent and Moran model simulations was almost perfect across the entire range of *ψ* (Fig. SI B_2-SI B_4). Only for very small *ψ*were slight deviations observed.

These results highlight two important points. First and reassuringly these results indicate that for large initial population sizes *N* the extended Moran model can be described by its limiting ancestral process (i.e., the psi-coalescent). Second, simulating the population dynamics forward in time is computationally intensive and prohibitively slow. However, since the ancestral process accurately captures the forward-in-time population dynamics, the coalescent process can be used to simulate the process quickly and efficiently over a large parameter space and for large sample sizes.

## Supporting Figures

**Figure SI B_1.**
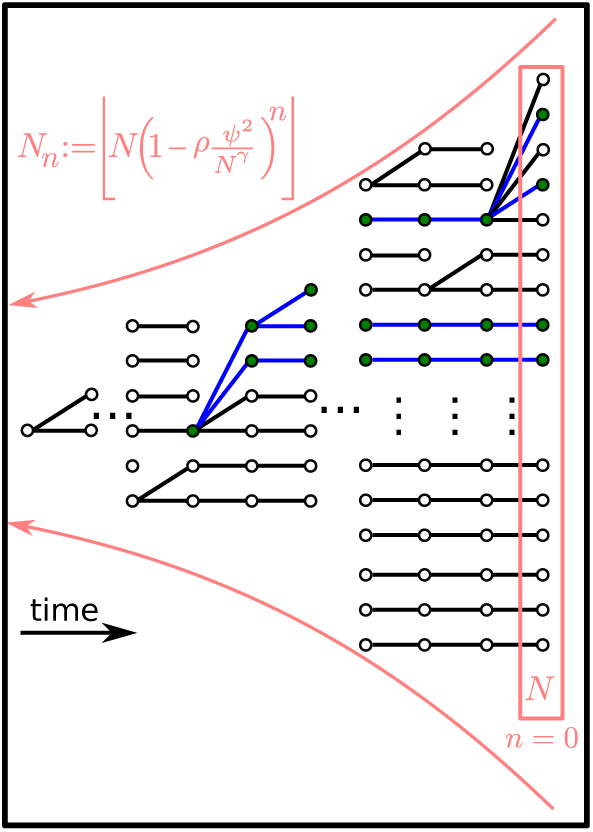
Illustration of genealogical relation in the extended Moran model with exponential growth. Circles denote individuals; time increases from left to right with the rightmost individuals constituting the present population. Green circles denote sampled individuals. Connecting lines between circles show the genealogical relationship between individuals; the genealogy of the sampled individuals is highlighted with blue. Pink lines illustrate that population size increases (forward in time) exponentially with rate *ρ*.

**Figure SI B_2.**
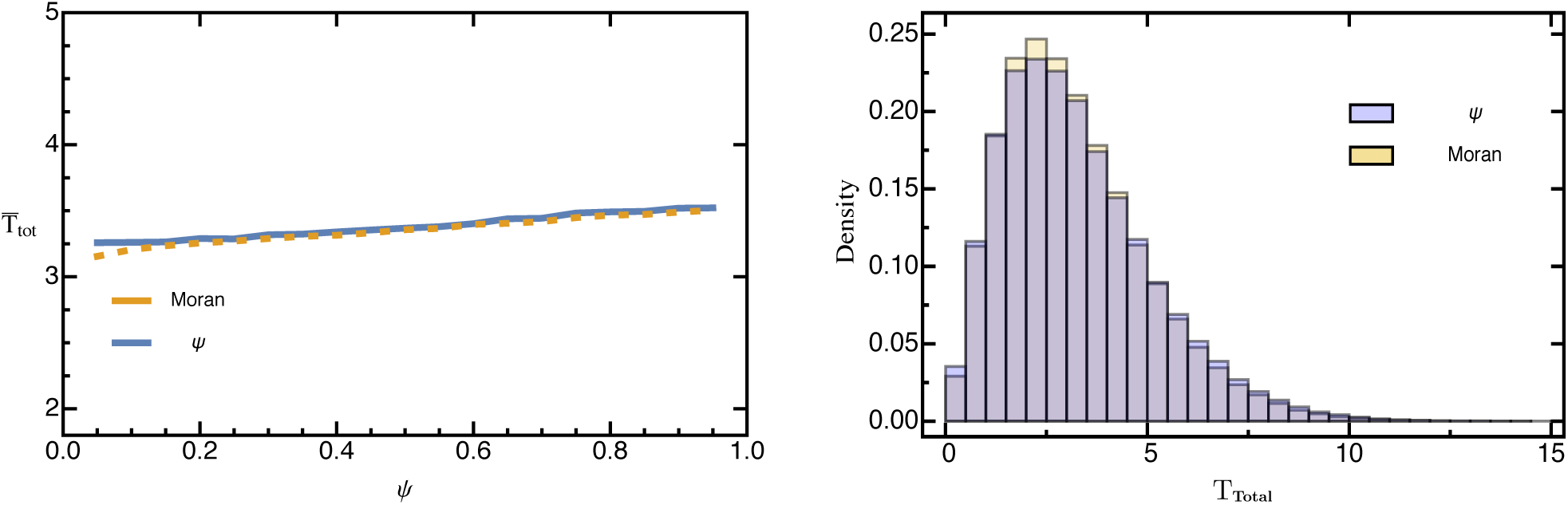
The left figure shows a comparison of 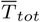 based on simulated trajectories of 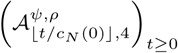 and 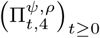 for ρ = 1, *γ* = 1.5, and different values of *ψ*. The right figure depicts the corresponding ns of 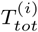 for *ψ* = 0.1. Averages were taken over 100,000 replicates. Other empirical distributio parameters: *N* = 100,000.

**Figure SI B_3.**
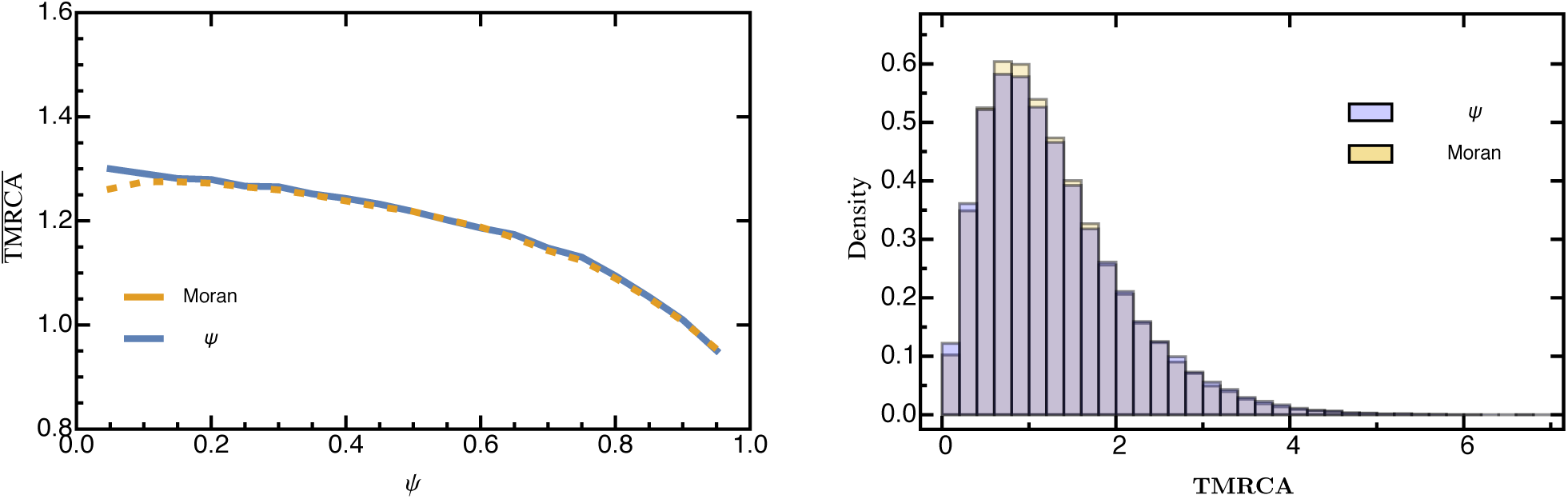
The left figure shows a comparison of *T̅_MRCA_* based on simulated trajectories of 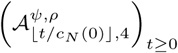 and 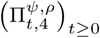 for *ρ* = 0.1, *γ* = 1.5, and different values of *ψ*. The right figure depicts the corresponding empirical distributions of 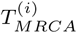 for *ψ* = 0.1. Averages were taken over 100,000 replicates. Other parameters: *N* = 100,000

**Figure SI B_4.**
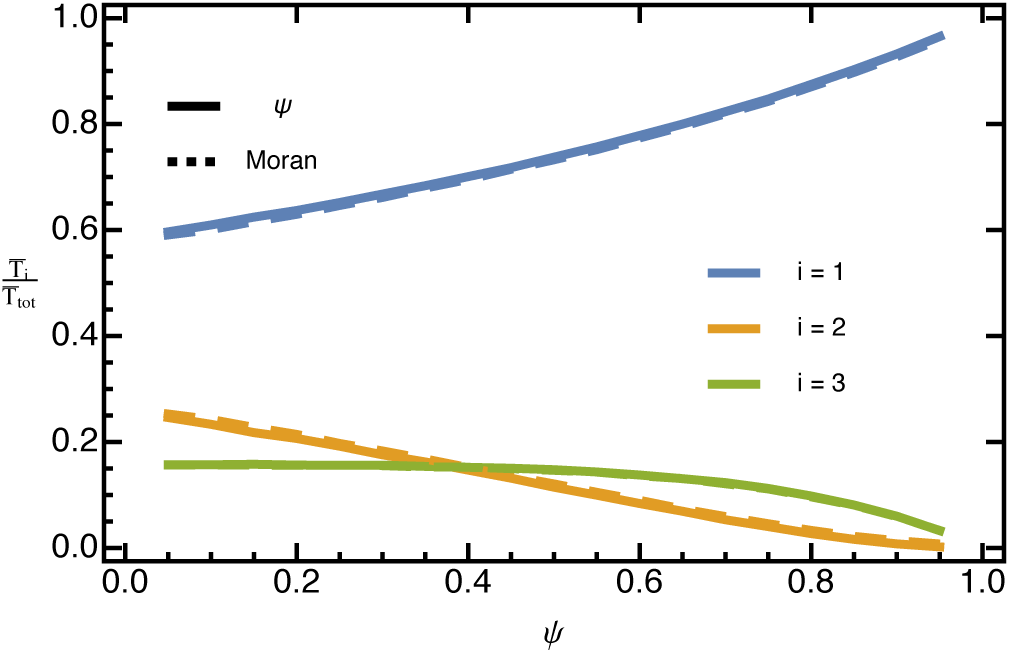
A comparison of 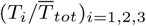 based on simulated trajectories of 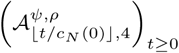 and 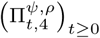 for *ρ* = 0.1, *γ* = 1.5, and different values of *ψ*. Averages were taken over 100,000 replicates. Other parameters: *N* = 100,000.

**Figure SI C_1.**
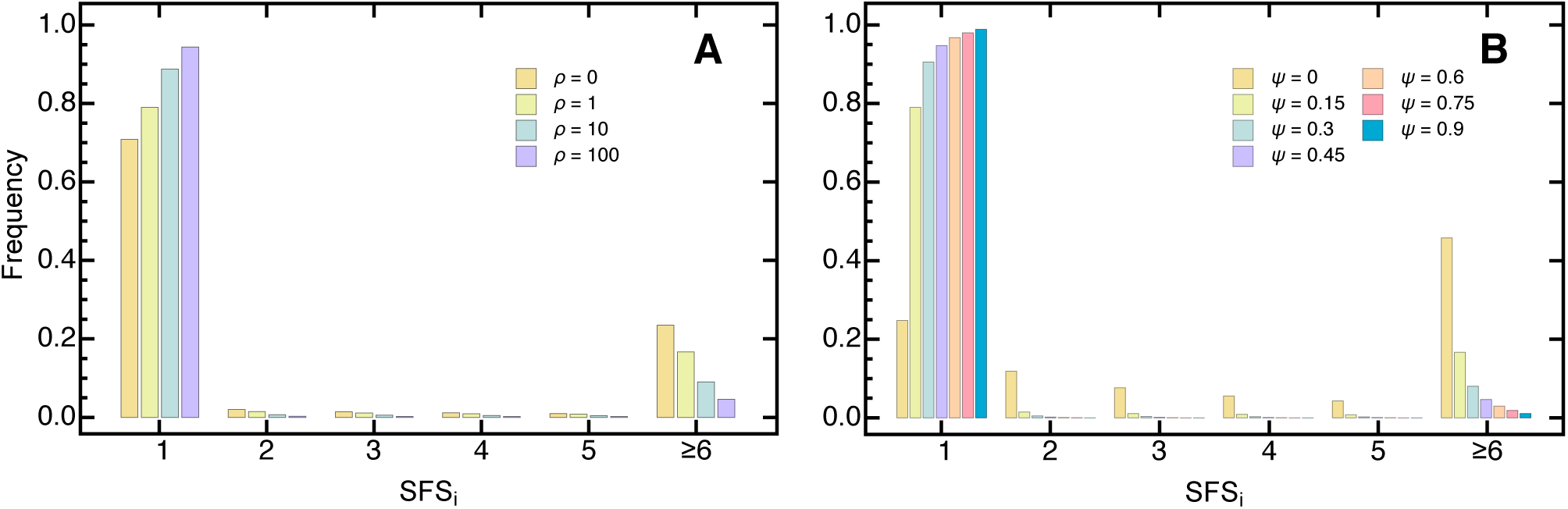
The normalized expected (lumped) SFS for the psi-coalescent for an exponentially growing population (eq. 18) with sample size *k* = 100 (**A**) for different values of *ρ* and fixed *ψ* = 0.15 and (**B**) for different values of *ψ* and fixed *ρ* = 1. The sixth entry in the SFS contains the aggregate of the higher frequency classes.

**Figure SI C_2.**
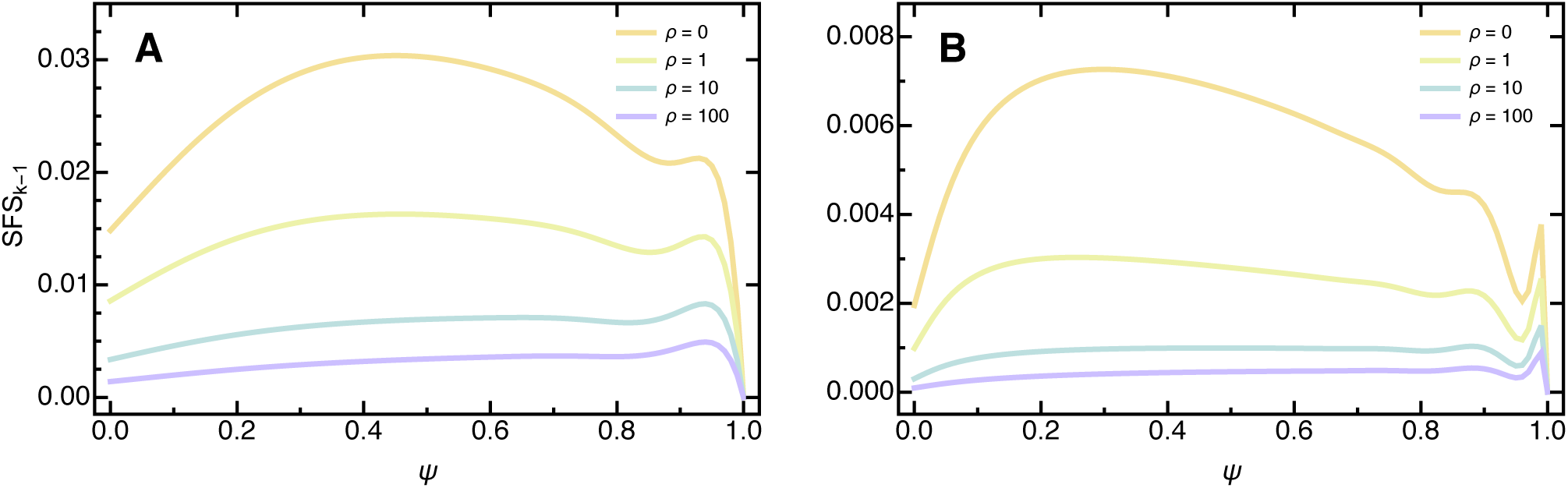
The normalized expected last entry of the SFS (eq. 18) as a function of *ψ* for various values of *ρ* with sample size (**A**) *k* = 20 and (**B**) *k* = 100.

**Figure SI C_3.**
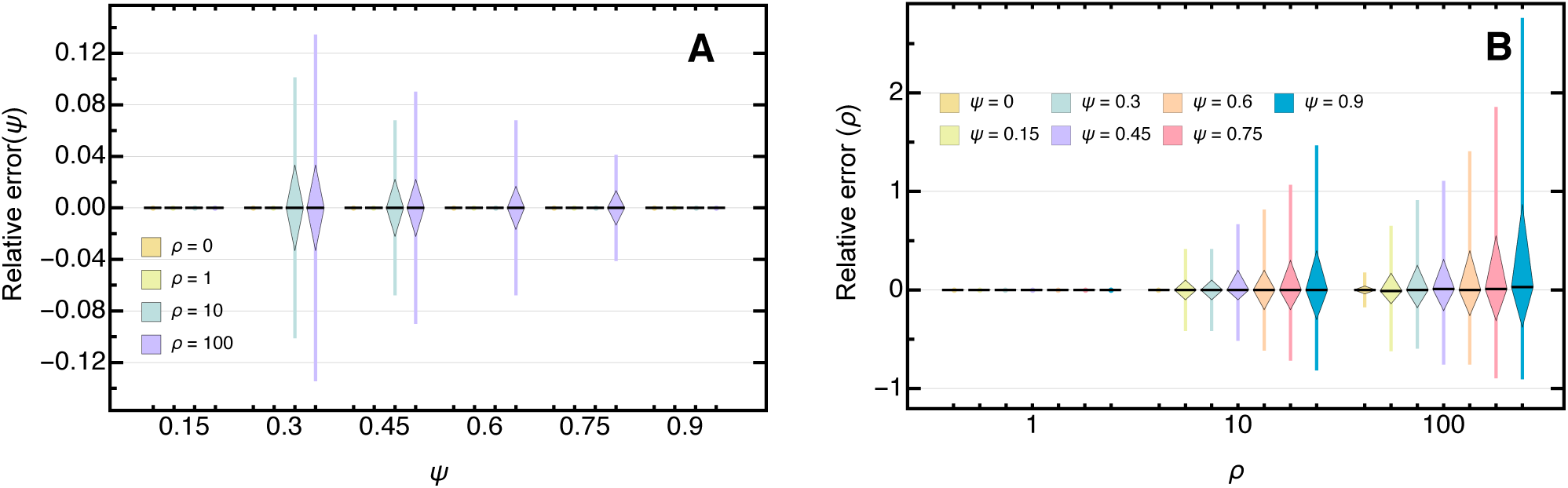
Boxplot of the relative error of the maximum likelihood estimate from the true (**A**) *ψ* and (**B**) *ρ* for 10,000 data sets assuming independent sites with *k* = 100 and *θ* (eq. 46) with *s* = 10,000. Boxes represent the interquartile range (i.e., the 50% C.I.) and whiskers extend to the highest/lowest data point within the box *±*1.5 times the interquartile range.

**Figure SI C_4.**
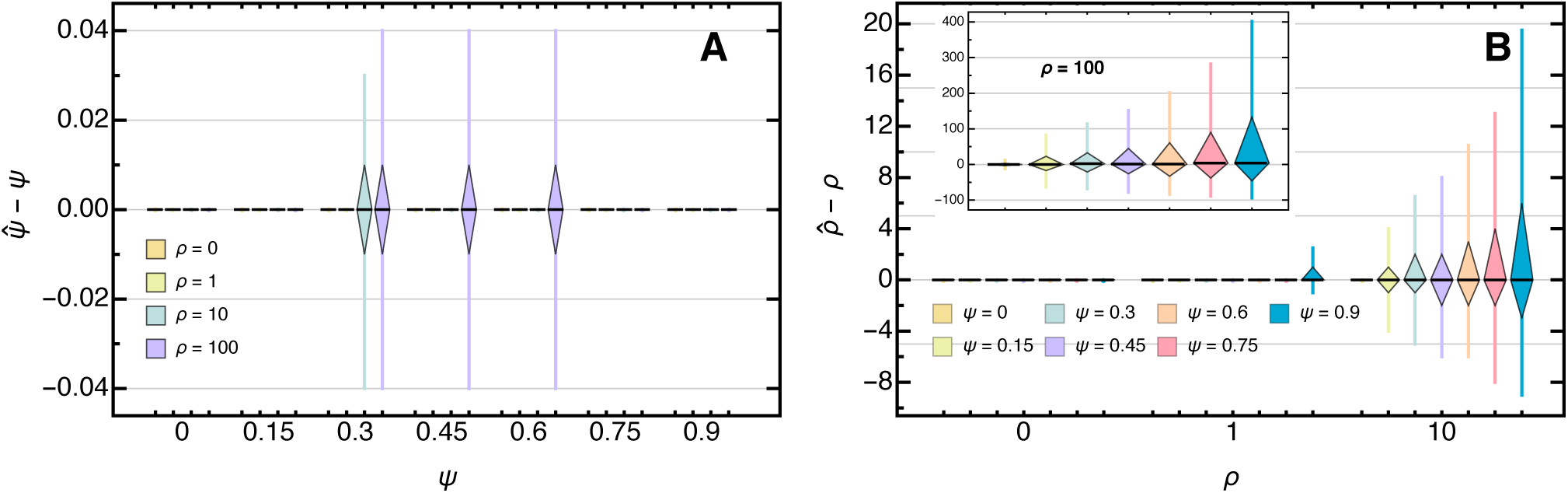
Boxplot of the deviation of the maximum likelihood estimate from the true (**A**) *ψ* and (**B**) *ρ* for 10,000 data sets assuming independent sites with *k* = 200 and and *θ* (eq. 46) with *s* = 10,000. Boxes represent the interquartile range (i.e., the 50% C.I.) and whiskers extend to the highest/lowest data point within the box *±*1.5 times the interquartile range. As the number of segregating sites *s* increases the variance of the estimator decreases and approaches its true underlying value.

**Figure SI C_5.**
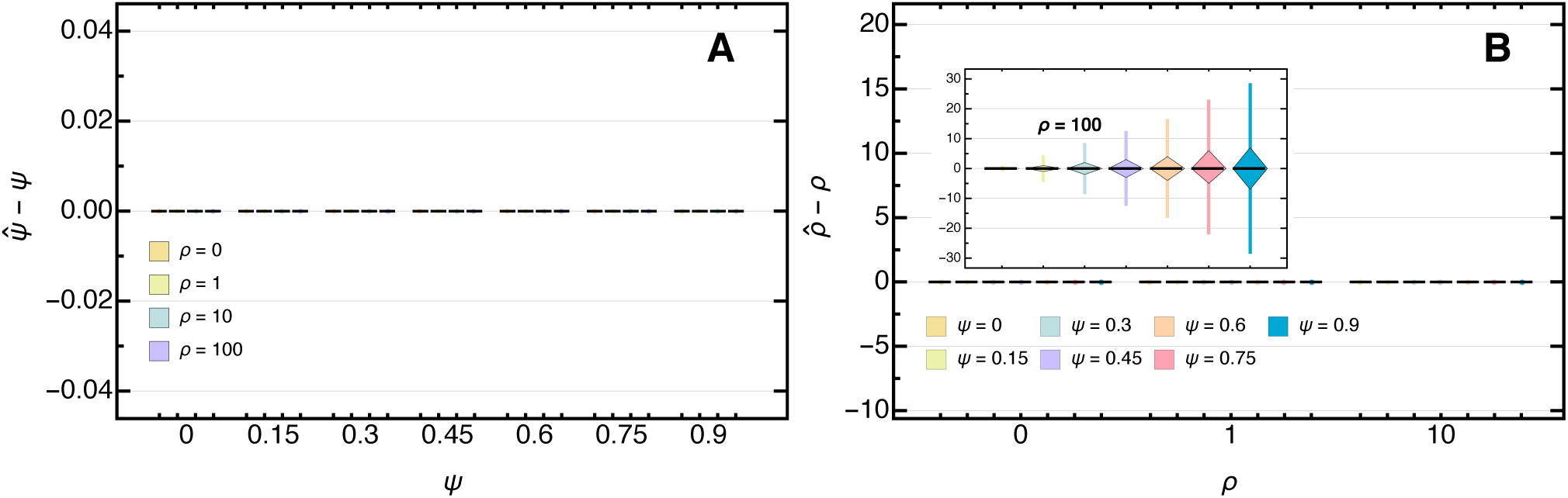
Boxplot of the deviation of the maximum likelihood estimate from the true (**A**) *ψ* and (**B**) *ρ* for 10,000 data sets assuming independent sites with *k* = 200 and *θ* (eq. 46) with *s* = 1,000,000. Boxes represent the interquartile range (i.e., the 50% C.I.) and whiskers extend to the highest/lowest data point within the box ±1.5 times the interquartile range. As the number of segregating sites *s* increases the variance of the estimator decreases and approaches its true underlying value.

**Figure SI C_6.**
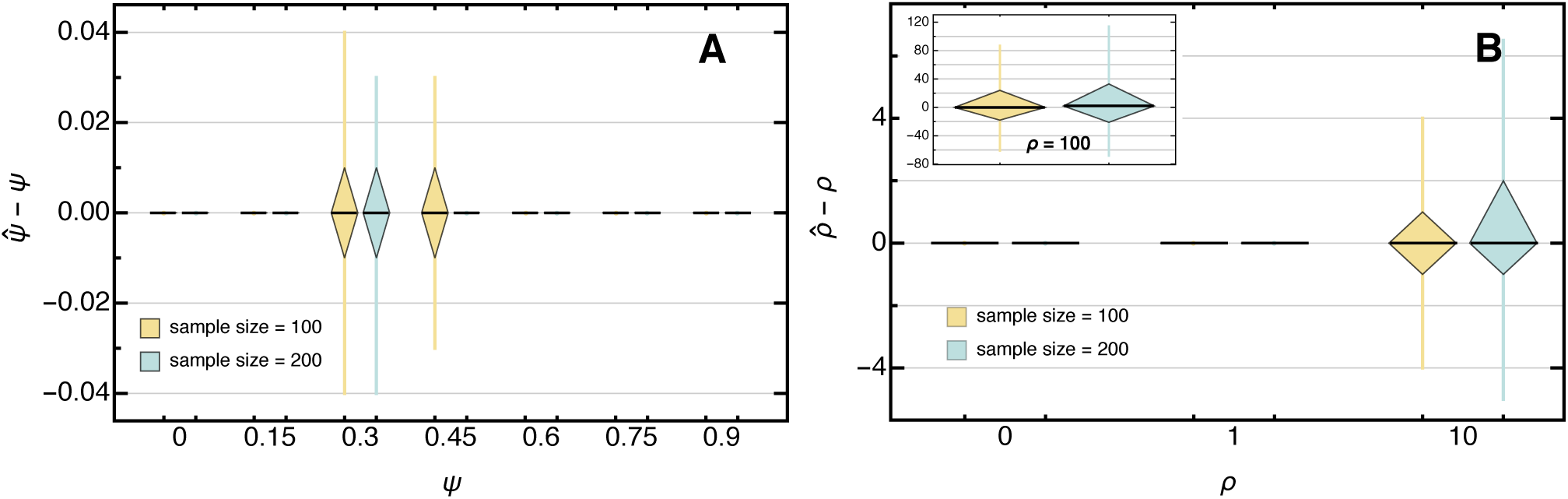
Boxplot of the deviation of the maximum likelihood estimate from the true (**A**) *ψ* and fixed *ρ* = 10, and (**B**) *ρ* and fixed *ψ* = 0.3 for 10,000 data sets assuming independent sites for different sample sizes *k* and *θ* (eq. 46) with *s* = 10,000. Boxes represent the interquartile range (i.e., the 50% C.I.) and whiskers extend to the highest/lowest data point within the box *±*1.5 times the interquartile range.

**Figure SI C_7.**
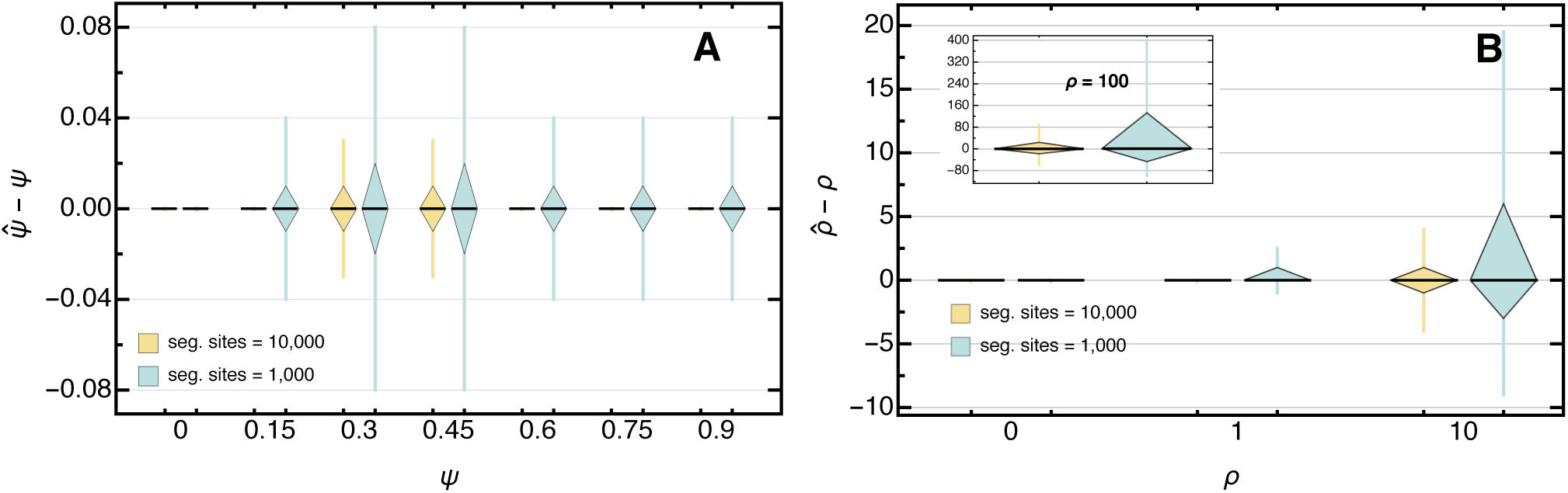
Boxplot of the deviation of the maximum likelihood estimate from the true (**A**) *ψ* and fixed *ρ* = 10, and (**B**) *ρ* and fixed *ψ* = 0.3 for 10,000 data sets assuming independent sites for different segregating sites *s* with *k* = 100. Boxes represent the interquartile range (i.e., the 50% C.I.) and whiskers extend to the highest/lowest data point within the box *±*1.5 times the interquartile range.

**Figure SI C_8.**
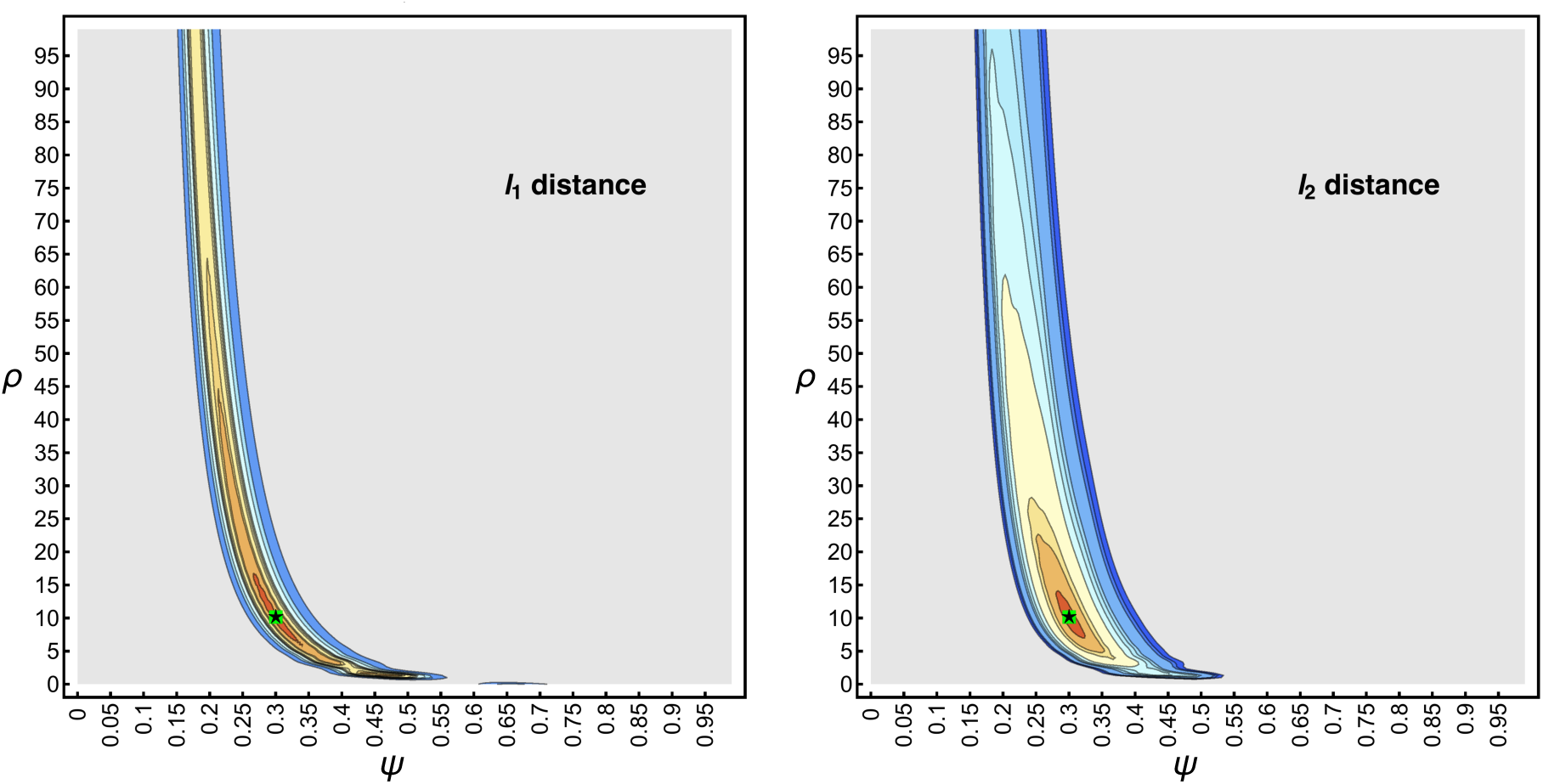
Surface of the *l*_1_ (left) and the *l*_2_ (right) distance of the idealized SFS with *k* = 100, *ψ* = 0.3, *ρ* = 10 and *s* = 10,000. Contours show the 0.95, 0.9675, 0.975, 0.99, 0.99225, 0.9945, 0.99675, 0.999, 0.99945 and 0.9999 quantiles. Distances below the 0.95 quantile are uniformly colored in gray. The green square shows the true *ψ* and *ρ*. The black star shows the minimum distance estimates 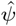and 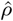.

**Figure SI C_9.**
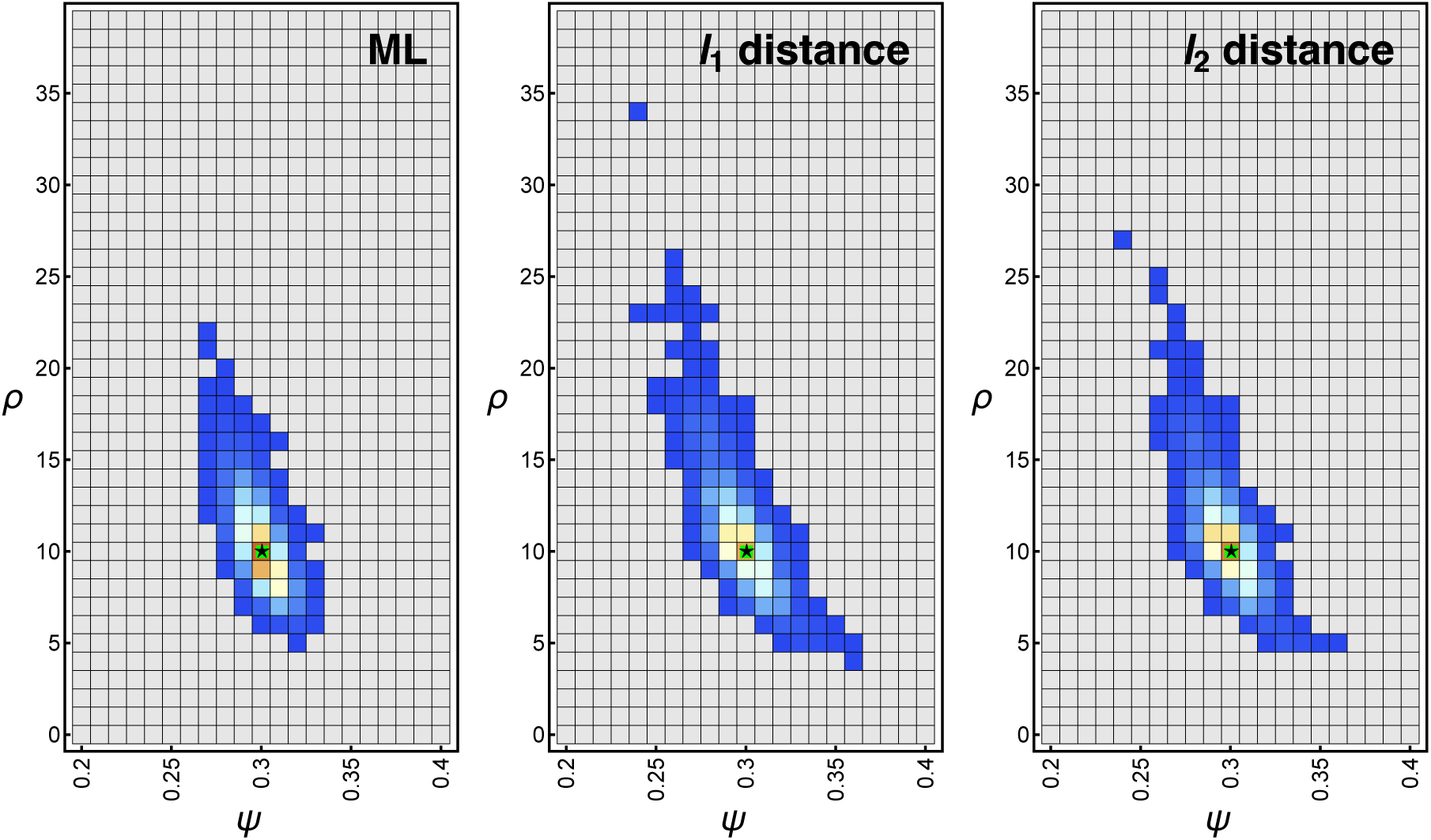
Heatplot of the frequency of the maximum likelihood estimates (left), the *l*_1_ distance (middle), and the *l*_2_ distance (right) for 10,000 data sets assuming independent sites with *k* = 100, *ψ* = 0.3, *ρ* = 10 and *s* = 10,000. Counts increase from blue to red with grey squares showing zero counts. The green square shows the true *ψ* and *ρ*. The black star shows the median (and mean) of the maximum likelihood respectively the minimum distance estimates 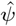and 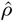.

**Figure SI C_10.**
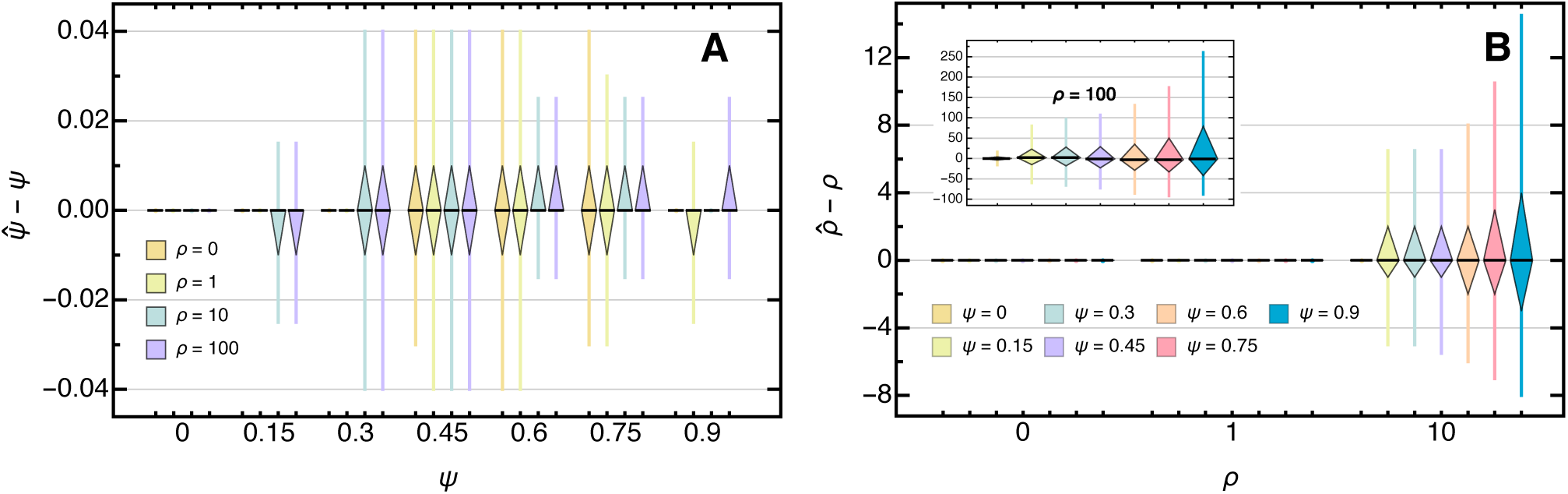
Boxplot of the deviation of the L1-distance estimate from the true (**A**) *ψ* and fixed *ρ* = 10, and (**B**) *ρ* and fixed *ψ* = 0.3 for 10,000 data sets assuming independent sites with *k* = 100 and *θ* (eq. 46) with *s* = 10,000. Boxes represent the interquartile range (i.e., the 50% C.I.) and whiskers extend to the highest/lowest data point within the box *±*1.5 times the interquartile range.

**Figure SI C_11.**
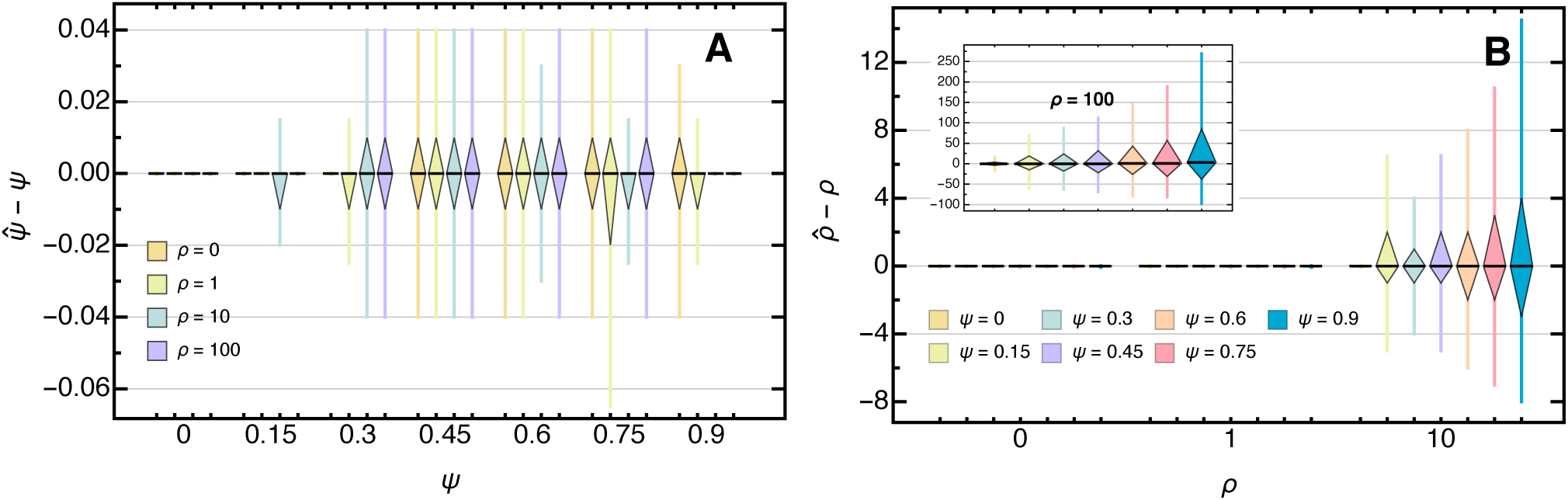
Boxplot of the deviation of the L2-distance estimate from the true (**A**) *ψ* and fixed *ρ* = 10, and (**B**) *ρ* and fixed *ψ* = 0.3 for 10,000 data sets assuming independent sites with *k* = 100 and *θ* (eq. 46) with *s* = 10,000‥ Boxes represent the interquartile range (i.e., the 50% C.I.) and whiskers extend to the highest/lowest data point within the box *±*1.5 times the interquartile range.

**Figure SI C_12.**
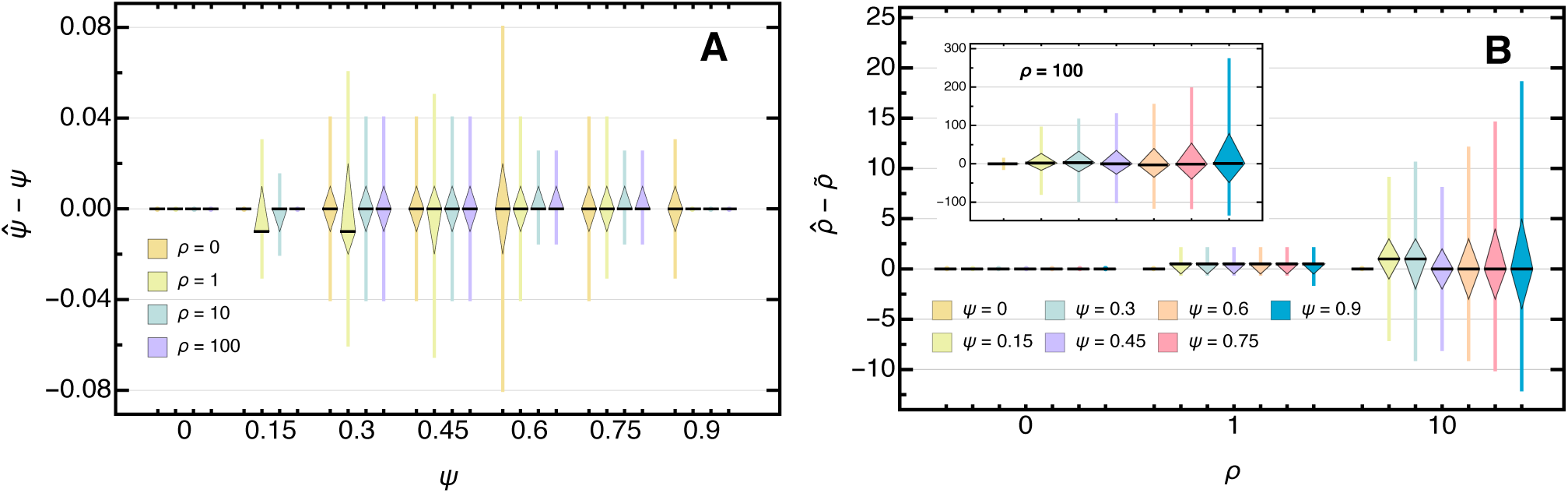
Boxplot of the deviation of the *l*_1_ distance estimate from the true (**A**) *ψ* and (**B**) *ρ* for 10,000 whole-genome data sets with *e* = 100*, k* = 100, *γ* = 1.5, and *θ* (eq. 46) with *s* = 1,000. Boxes represent the interquartile range (i.e., the 50% C.I.) and whiskers extend to the highest/lowest data point within the box *±*1.5 times the interquartile range.

**Figure SI C_13.**
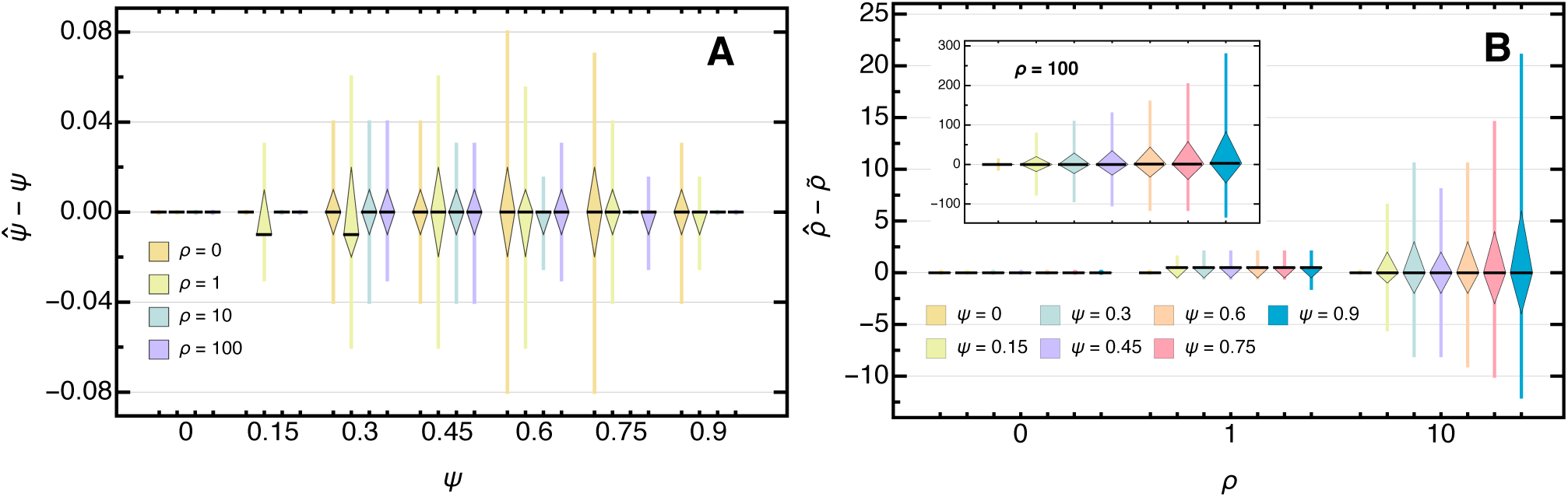
Boxplot of the deviation of the *l*_2_ distance estimate from the true (**A**) *ψ* and (**B**) *ρ* for 10,000 whole-genome data sets with *l* = 100*, k* = 100, *γ* = 1.5, and *θ* (eq. 46) with *s* = 1,000. Boxes represent the interquartile range (i.e., the 50% C.I.) and whiskers extend to the highest/lowest data point within the box *±*1.5 times the interquartile range.

**Figure SI C_14.**
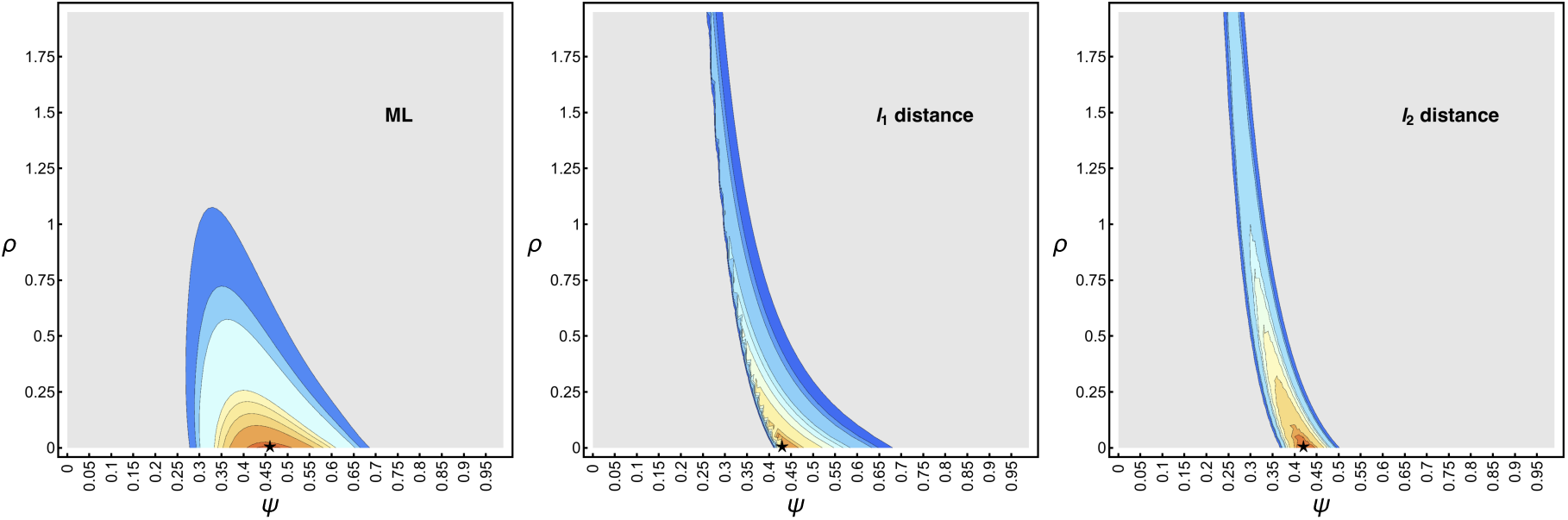
Likelihood (left), *l*_1_ distance, and *l*_2_ distance surface of the unfolded SFS (given the ML rooted tree) of the sardine mtDNA sequences with *k* = 106 and *s* = 81. Contours show the corresponding 0.95, 0.9675, 0.975, 0.99, 0.99225, 0.9945, 0.99675, 0.999, 0.99945 and 0.9999 quantiles. Likelihoods respectively distances below their corresponding 0.95 quantile are uniformly colored in gray. The black star shows the maximum likelihood respectively minimum distance estimates: 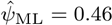 and 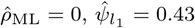 and 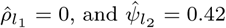 and 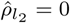.

## Supporting Tables

**Table SI D_1.**
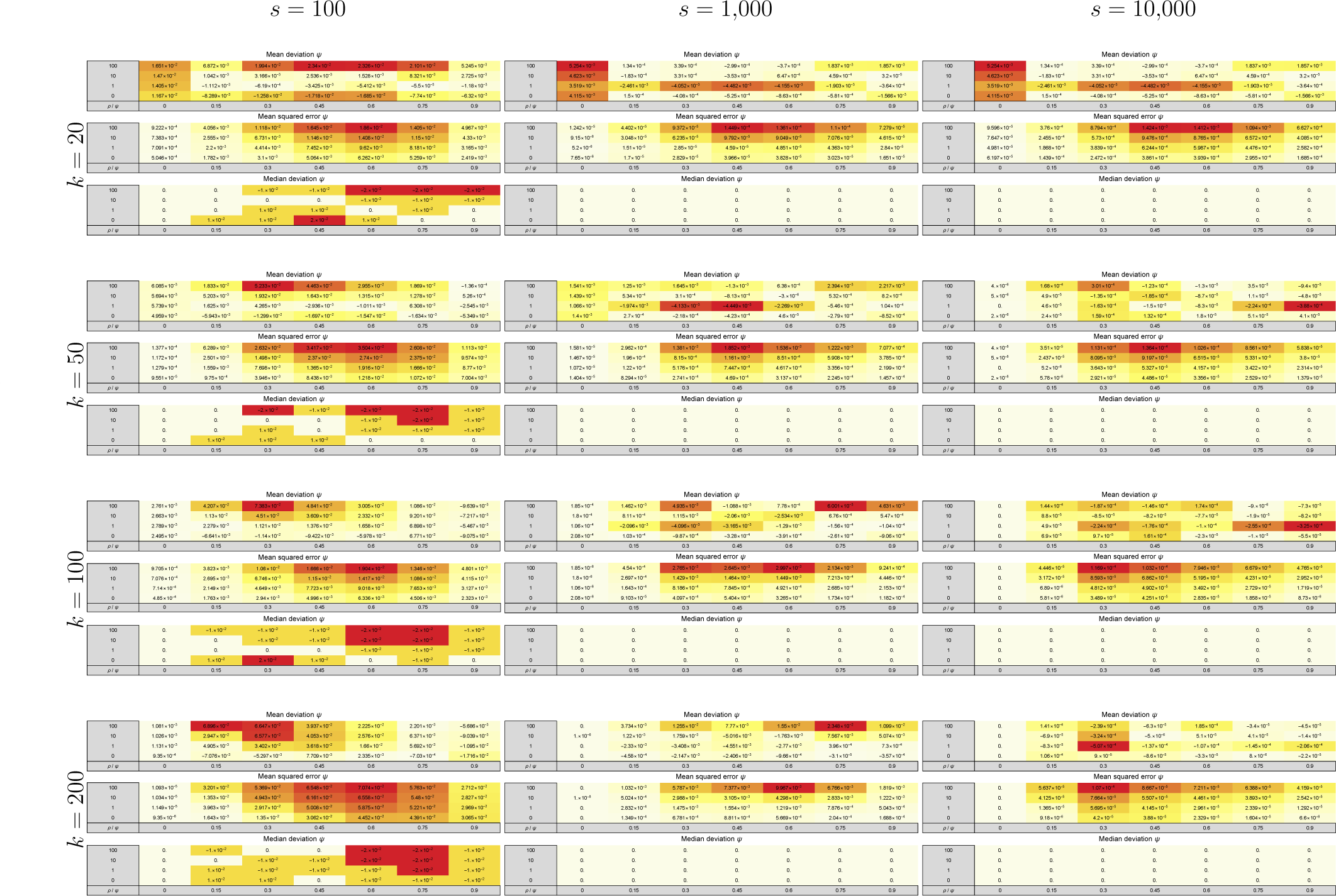
Overview of the (marginal) accuracy 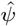 when assuming independent sites. Each cell shows the mean difference (first row), the mean squared error (second row) and the median difference (third row) of 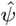 calculated over 10,000 data sets assuming independent sites. Colors within each sub-table range from light yellow to dark red and scale between the minimal and the maximal absolute value to aid interpretation.

**Table SI D_2.**
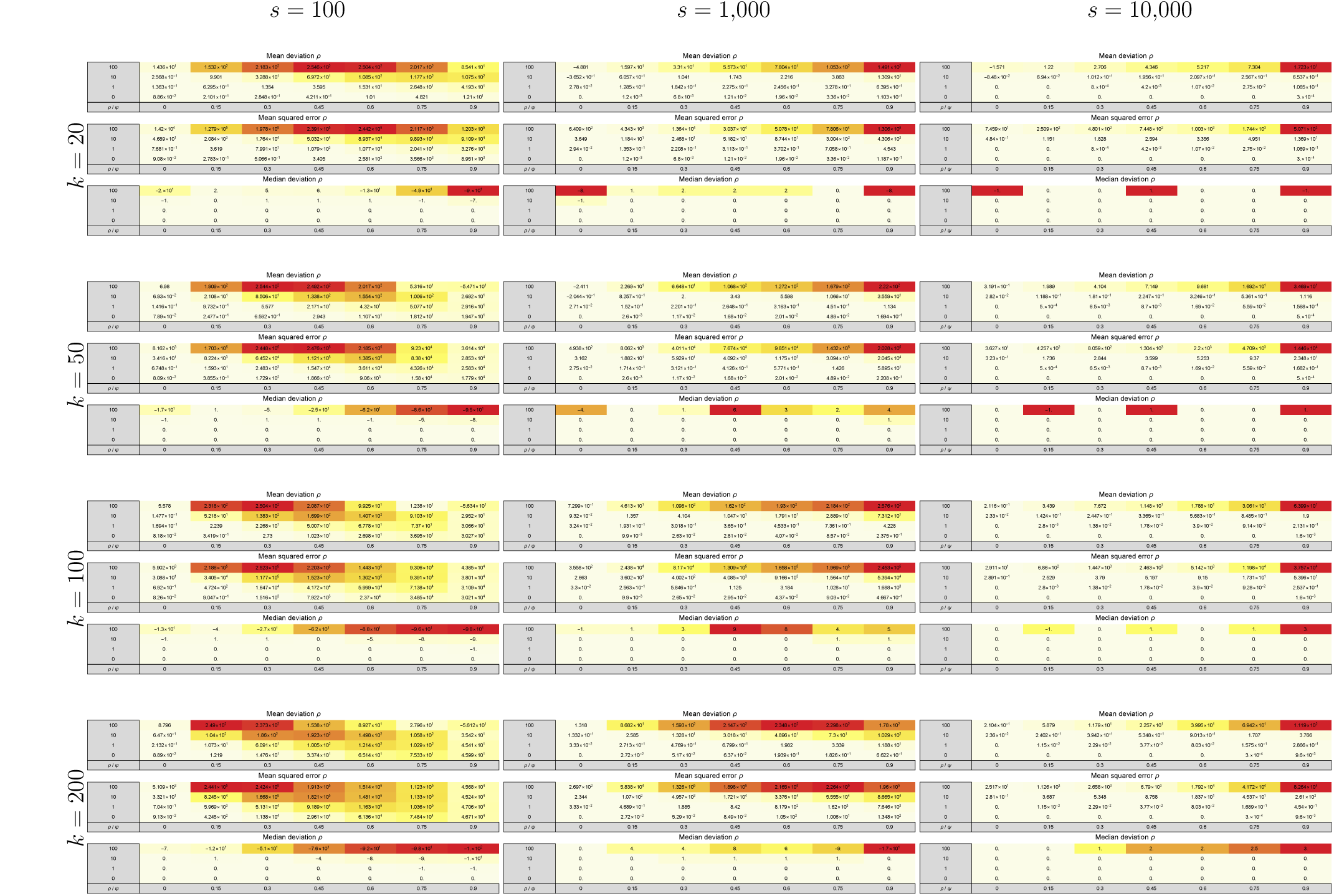
Overview of the (marginal) accuracy 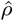 when assuming independent sites. Each cell shows the mean difference (first row), the mean squared error (second row) and the median difference (third row) of 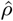 (right column) calculated over 10,000 data sets assuming independent sites. Colors within each sub-table range from light yellow to dark red and scale between the minimal and the maximal absolute value to aid interpretation.

**Table SI D_3.**
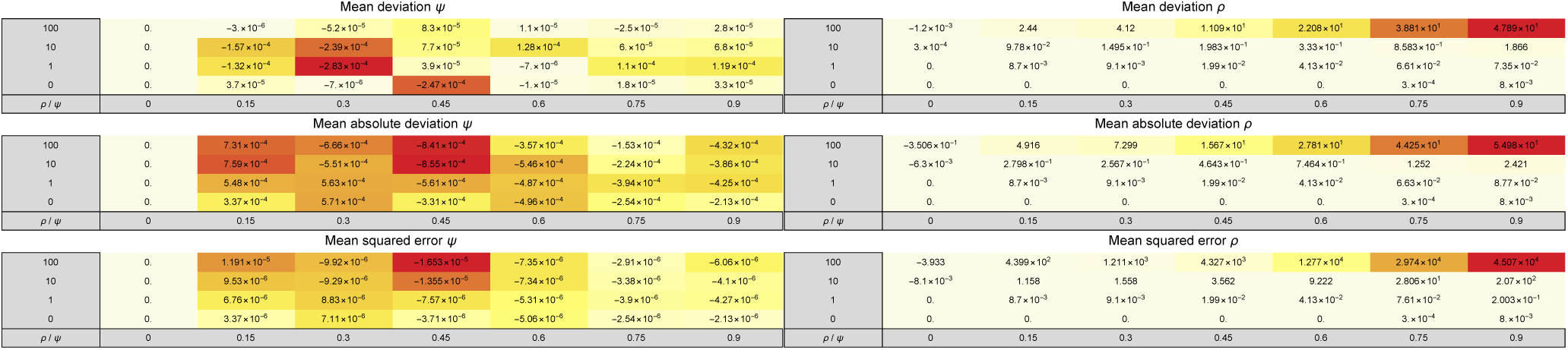
Comparison of the (marginal) accuracy of 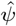 (left column) and 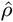 (right column) for *k* = 100 (i.e., the reference) and *k* = 200 when assuming independent sites. Each cell shows the difference of the absolute mean difference (|*M D*_*k*=100_|–|*M D*_*k*=200_|; first row), the difference of the mean absolute difference (*M AD*_*k*=100_ – *M AD*_*k*=200_; third row) and the difference of the mean squared error (*M SE*_*k*=100_ – *M SE*_*k*=200_; third row) each calculated over 10,000 data sets assuming independent sites and *θ* (eq. 46) with *s* = 10,000. Colors within each sub-table range from light yellow to dark red and scale between the minimal and the maximal absolute value to aid interpretation.

**Table SI D_4.**
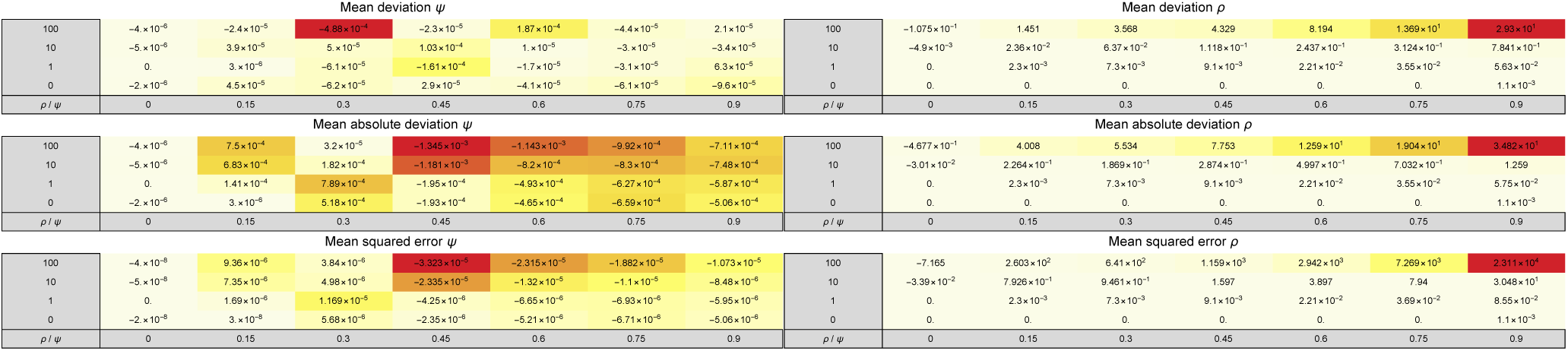
Comparison of the (marginal) accuracy of 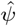 (left column) and 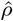 (right column) for *k* = 100 (i.e., the reference) and *k* = 50 when assuming independent sites. Each cell shows the difference of the absolute mean difference (|*M D*_*k*=100_| – |*M D*_*k*=50_|; first row), the difference of the mean absolute difference (*M AD*_*k*=100_ – *M AD*_*k*=50_; third row) and the difference of the mean squared error (*M SE*_*k*=100_ − *M SE*_*k*=50_; third row) each calculated over 10,000 data sets assuming independent sites and *θ* (eq. 46) with *s* = 10,000. Colors within each sub-table range from light yellow to dark red and scale between the minimal and the maximal absolute value to aid interpretation.

**Table SI D_5.**
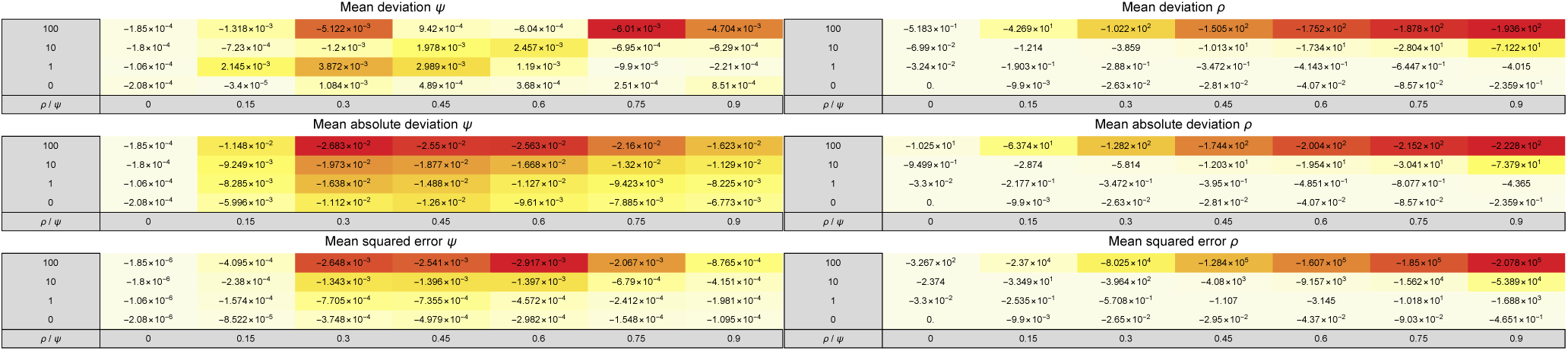
Comparison of the (marginal) accuracy of 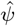(left column) and 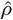 (right column) for *θ* (eq. 46) with *s* = 10,000 (i.e., the reference) and *s* = 1,000 when assuming independent sites. Each cell shows the difference of the absolute mean difference (|*M D*_*s*=10,000_|– |*M D*_*s*=1,000_|; first row), the difference of the mean absolute difference (*M AD*_*s*=10,000_–*M AD*_*s*=1,000_; third row) and the difference of the mean squared error (*M SE*_*s*=10,000_–*M SE*_*s*=1,000_; third row) each calculated over 10,000 data sets assuming independent sites and *k* = 100. Colors within each sub-table range from light yellow to dark red and scale between the minimal and the maximal absolute value to aid interpretation.

**Table SI D_6.**
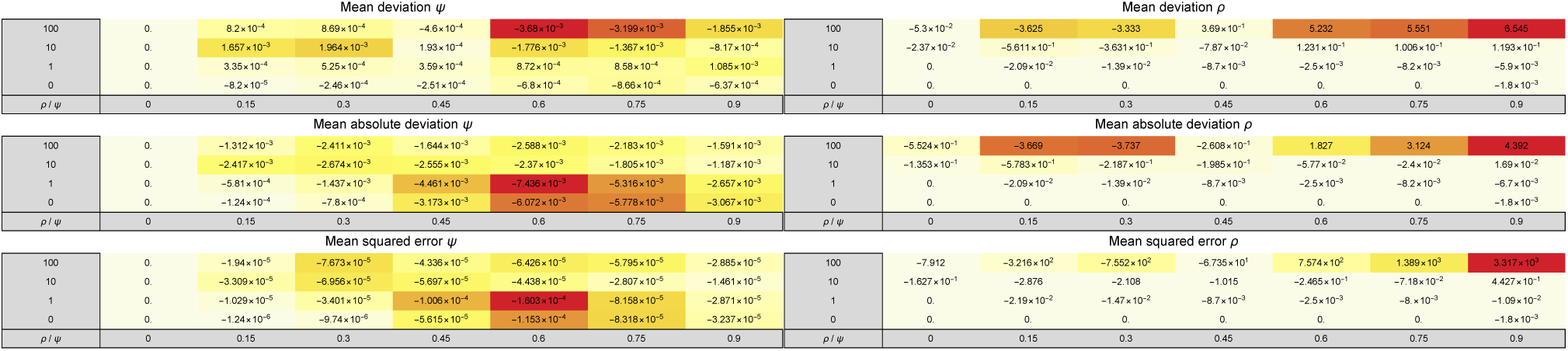
Comparison of the (marginal) accuracy of 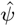 (left column) and 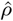 (right column) for the maximum likelihood-based estimate (i.e., the reference) and the L1-distance-based estimate when assuming independent sites. Each cell shows the difference of the absolute mean difference (|*M D*_*MLE*_| –*|M D*_*L*1_|; first row), the difference of the mean absolute difference (*M AD*_*MLE*_*M AD*_*L*1_; third row) and the difference of the mean squared error (*M SE*_*MLE*_*–M SE*_*L*1_; third row) each calculated over 10,000 data sets assuming independent sites, *k* = 100, and *θ* (eq. 46) with *s* = 10,000. Colors within each sub-table range from light yellow to dark red and scale between the minimal and the maximal absolute value to aid interpretation.

**Table SI D_7.**
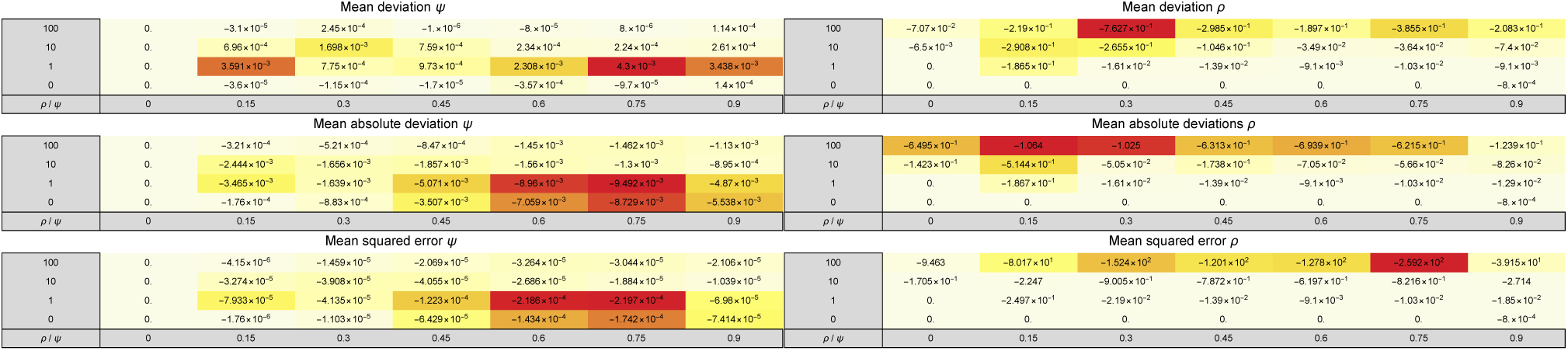
Comparison of the (marginal) accuracy of 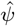 (left column) and 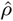 (right column) for the maximum likelihood-based estimate (i.e., the reference) and the L2-distance-based estimate when assuming independent sites. Each cell shows the difference of the absolute mean difference (|*M D*_*MLE*_|*–|M D*_*L*2_|; first row), the difference of the mean absolute difference (*M AD*_*MLE*_*M AD*_*L*2_; third row) and the difference of the mean squared error (*M SE*_*MLE*_*– M SE*_*L*2_; third row) each calculated over 10,000 data sets assuming independent sites, *k* = 100, and *θ* (eq. 46) with *s* = 10,000. Colors within each sub-table range from light yellow to dark red and scale between the minimal and the maximal absolute value to aid interpretation.

**Table SI D_8.**
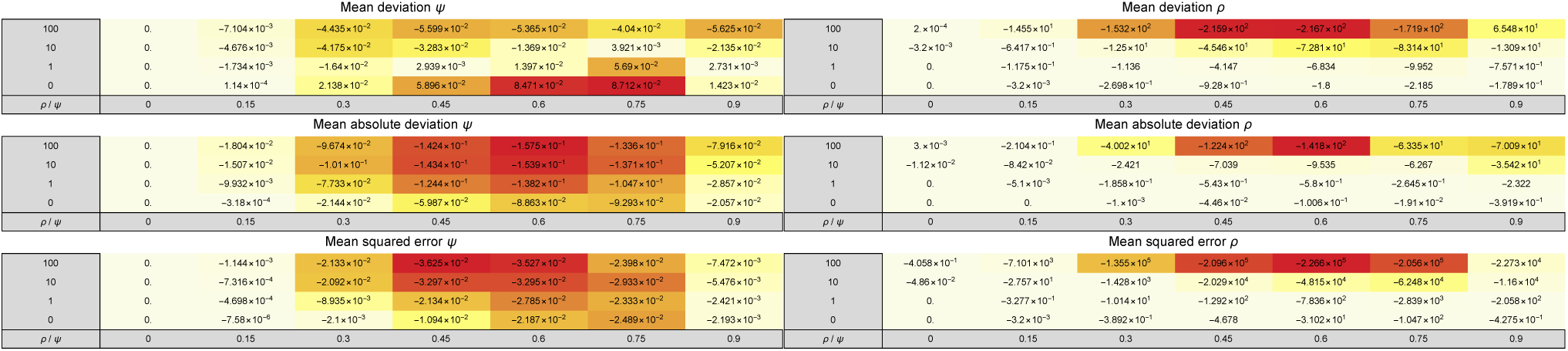
Comparison of the (marginal) accuracy of 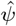 (left column) and 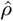 (right column) for the maximum likelihood estimate based on the full SFS (i.e., the reference) and the lumped SFS where the *i* = 5 entry in the SFS contains the aggregate of the higher frequency classes when assuming independent sites‥ Each cell shows the difference of the absolute mean difference (|*M D*_*i*=0_| –|*M D*_*i*=5_|; first row), the difference of the mean absolute difference (*M AD*_*i*=0_ − *M AD*_*i*=5_; third row) and the difference of the mean squared error (*M SE*_*i*=0_ − *M SE*_*i*=5_; third row) each calculated over 10,000 data sets assuming independent sites, *k* = 100, and *θ* (eq. 46) with *s* = 10,000. Colors within each sub-table range from light yellow to dark red and scale between the minimal and the maximal absolute value to aid interpretation.

**Table SI D_9.**
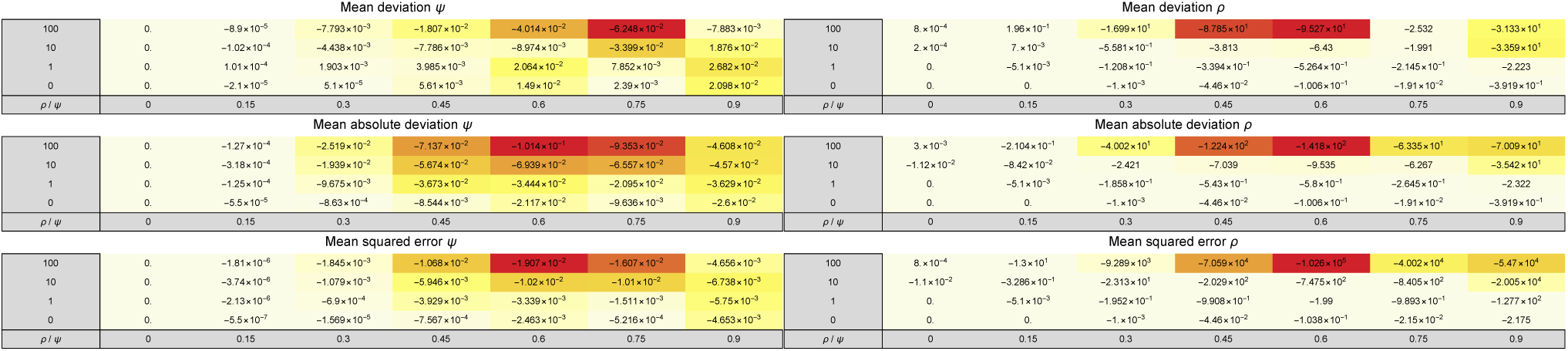
Comparison of the (marginal) accuracy of 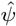 (left column) and 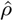 (right column) for the maximum likelihood estimate based on the full SFS (i.e., the reference) and the lumped SFS where the *i* = 15 entry in the SFS contains the aggregate of the higher frequency classes when assuming independent sites‥ Each cell shows the difference of the absolute mean difference (|*M D*_*i*=0_| – |*M D*_*i*=15_|; first row), the difference of the mean absolute difference (*M AD*_*i*=0_ − *M AD*_*i*=15_; third row) and the difference of the mean squared error (*M SE*_*i*=0_ − *M SE*_*i*=15_; third row) each calculated over 10,000 data sets assuming independent sites, *k* = 100, and *θ* (eq. 46) with *s* = 10,000. Colors within each sub-table range from light yellow to dark red and scale between the minimal and the maximal absolute value to aid interpretation.

**Table SI D_10.**
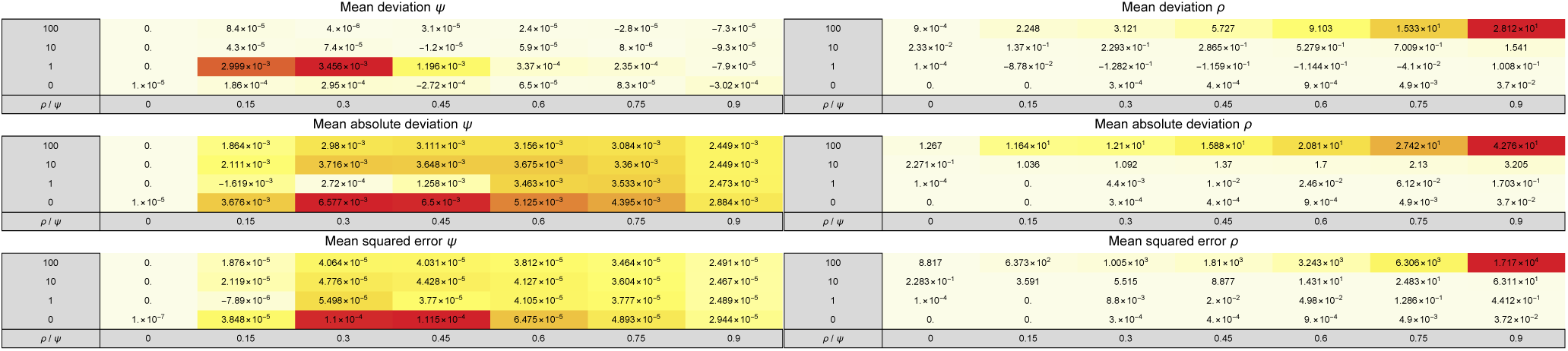
Comparison of the (marginal) accuracy of 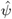(left column) and 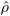(right column) for the maximum likelihood estimate based for *l* = 100 (i.e., the reference) and *l* = 1,000. Each cell shows the difference of the absolute mean difference (|*M D*_*l*=100_|–|*M D*_*l*=1,000_|; first row), the difference of the mean absolute difference (*M AD*_*l*=100_ – *M AD*_*l*=1,000_; third row) and the difference of the mean squared error (*M SE*_*l*=100_ –*M SE*_*l*=1,000_; third row) each calculated over 10,000 data sets assuming independent sites, *k* = 100, and *θ* (eq. 46) with *s* = 1,000 for *l* = 100 and *s* = 100 for *l* = 1,000. Colors within each sub-table range from light yellow to dark red and scale between the minimal and the maximal absolute value to aid interpretation.

**Table SI D_11.**
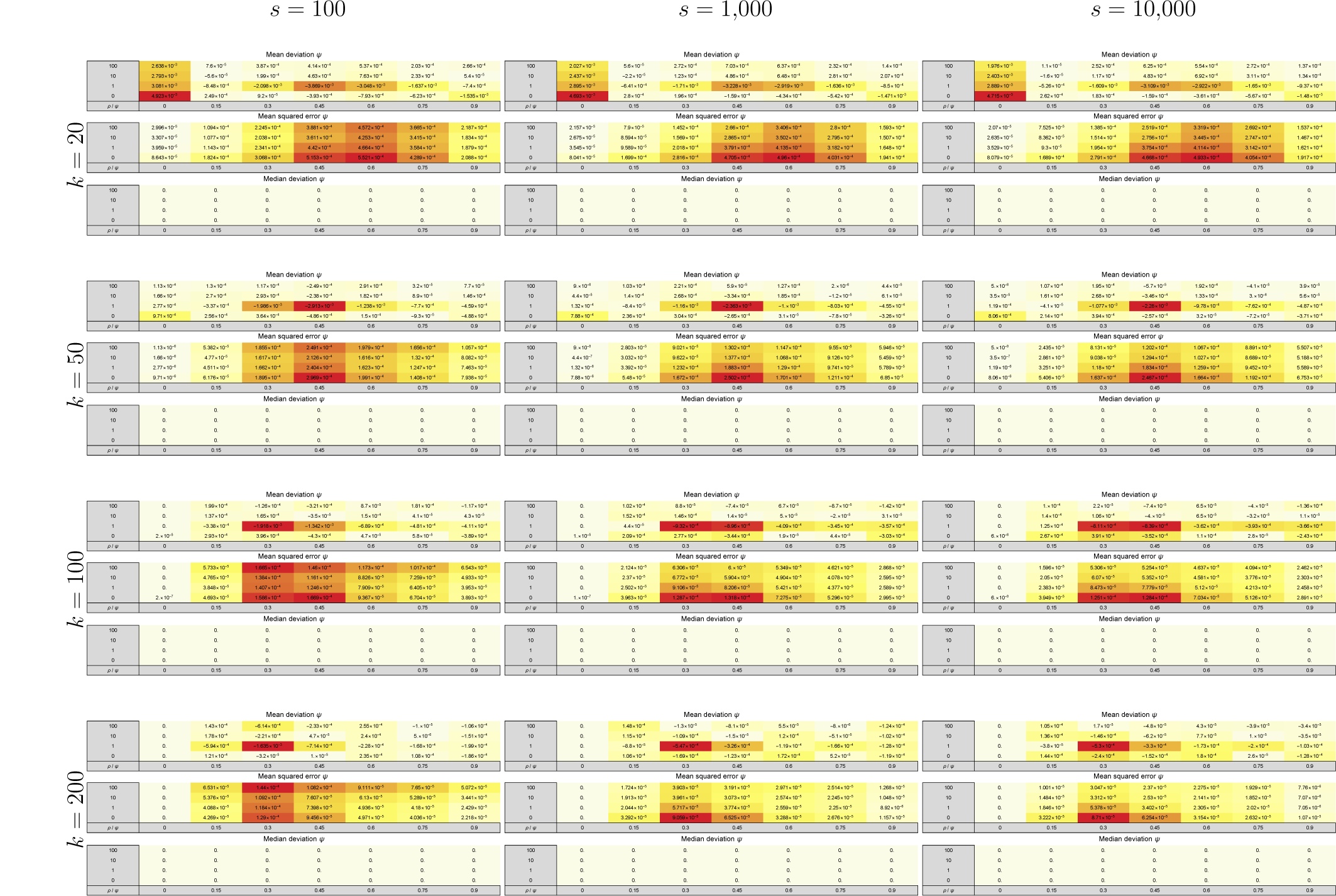
Overview of the (marginal) accuracy 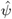 estimated from whole-genome simulations with *l* = 100. Each cell shows the mean difference (first row), the mean squared error (second row) and the median difference (third row) of 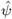 calculated over 10,000 data sets assuming independent sites. Colors within each sub-table range from light yellow to dark red and scale between the minimal and the maximal absolute value to aid interpretation.

**Table SI D_12.**
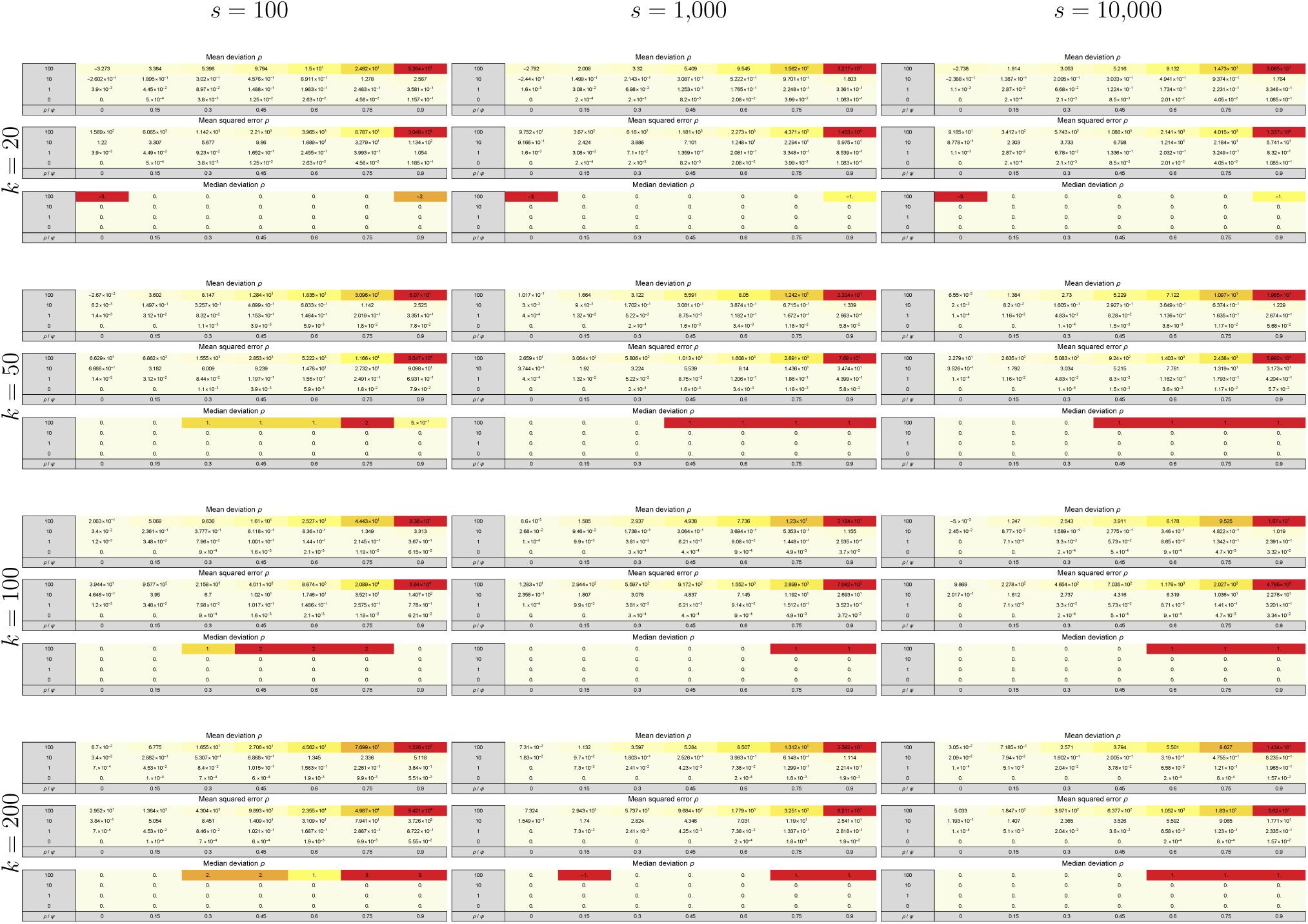
Overview of the (marginal) accuracy 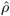 estimated from whole-genome simulations with *l* = 100. Each cell shows the mean difference (first row), the mean squared error (second row) and the median difference (third row) of 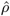 calculated over 10,000 data sets assuming independent sites. Colors within each sub-table range from light yellow to dark red and scale between the minimal and the maximal absolute value to aid interpretation.

**Table SI D_13.**
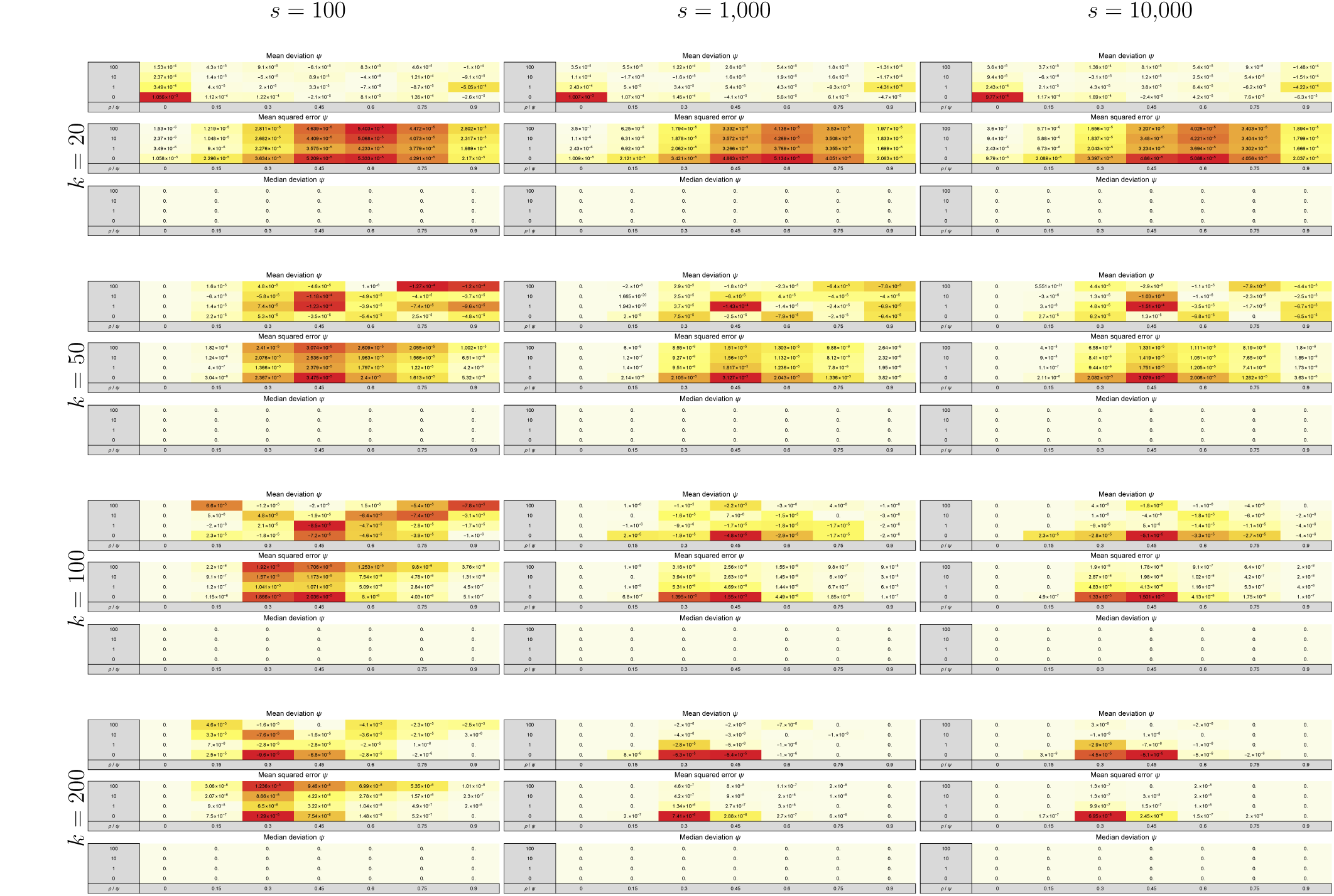
Overview of the (marginal) accuracy 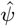 estimated from whole-genome simulations with *l* = 1,000. Each cell shows the mean difference (first row), the mean squared error (second row) and the median difference (third row) of 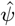 calculated over 10,000 data sets assuming independent sites. Colors within each sub-table range from light yellow to dark red and scale between the minimal and the maximal absolute value to aid interpretation.

**Table SI D_14.**
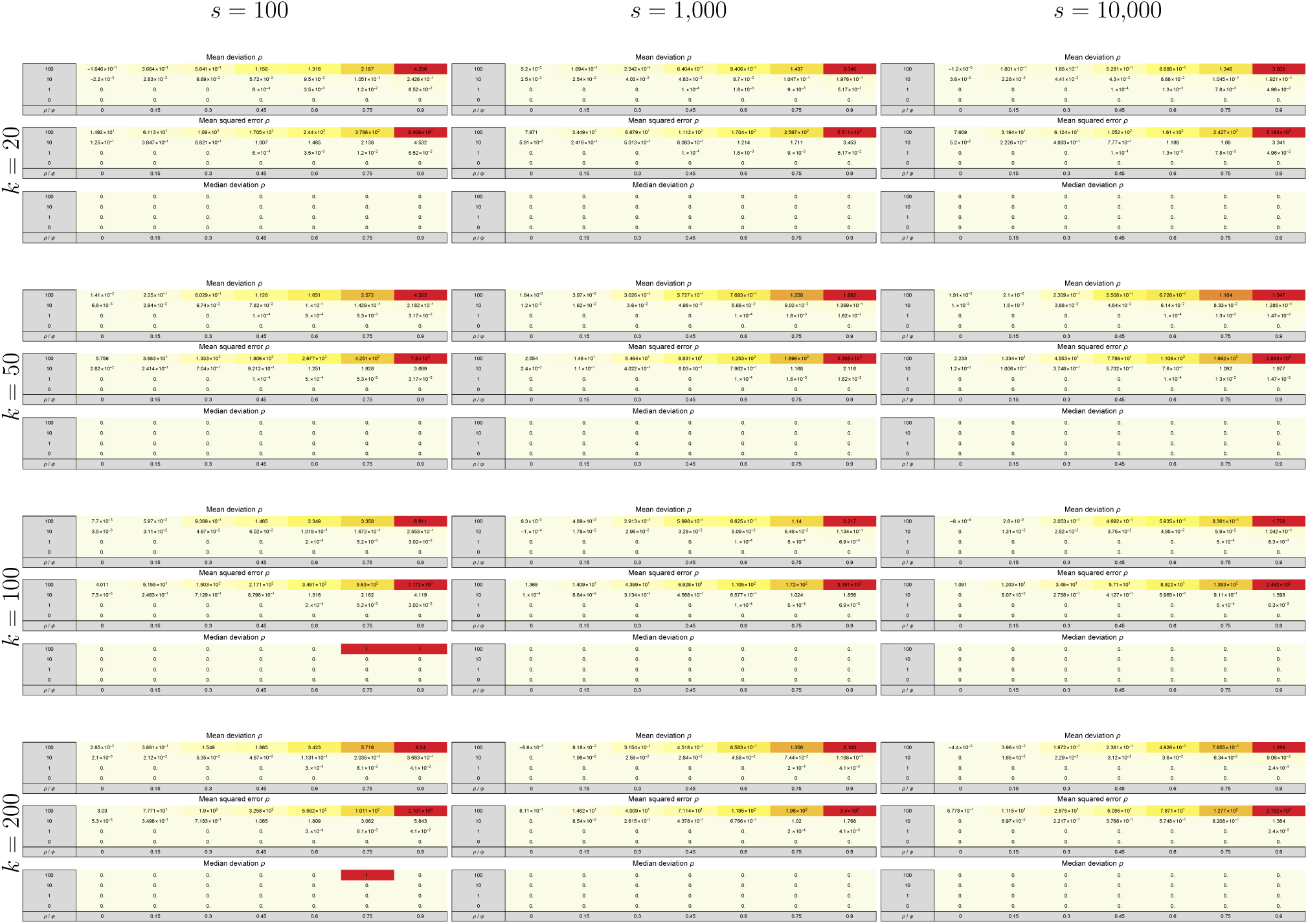
Overview of the (marginal) accuracy 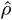 estimated from whole-genome simulations with *l* = 1,000. Each cell shows the mean difference (first row), the mean squared error (second row) and the median difference (third row) of 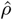 calculated over 10,000 data sets assuming independent sites. Colors within each sub-table range from light yellow to dark red and scale between the minimal and the maximal absolute value to aid interpretation.

## Supporting Files

File E1: SFS-Sardine-NiwaEtAl2016.txt

